# Nanoscale domains govern local diffusion and aging within FUS condensates

**DOI:** 10.1101/2024.04.01.587651

**Authors:** Guoming Gao, Emily R. Sumrall, Nils G. Walter

## Abstract

Biomolecular condensates regulate cellular physiology by sequestering and processing RNAs and proteins, yet how these processes are locally tuned within condensates remains unclear. Moreover, in neurodegenerative diseases such as amyotrophic lateral sclerosis (ALS), condensates undergo liquid-to-solid phase transitions, but capturing early intermediates in this process has been challenging. Here, we present a surface multi-tethering approach to achieve intra-condensate single-molecule tracking of fluorescently labeled RNA and protein molecules within liquid-like condensates. Using RNA-binding protein Fused in Sarcoma (FUS) as a model for condensates implicated in ALS, we discover that RNA and protein diffusion is confined within distinct nanometer-scale domains, or nanodomains, which exhibit unique connectivity and chemical environments. During condensate aging, these nanodomains reposition, facilitating FUS fibrilization at the condensate surface, a transition enhanced by FDA-approved ALS drugs. Our findings demonstrate that nanodomain formation governs condensate function by modulating biomolecule sequestration and percolation, offering insights into condensate aging and disease-related transitions.

## Main

Biomolecular condensates play a fundamental role in cellular physiology by regulating biochemical reactions through the phase separation of RNAs and proteins^1–4^. Among them, ribonucleoprotein (RNP) condensates are critical for RNA metabolism^5^ under both physiological and pathological conditions^4^. These condensates selectively recruit RNA molecules and their processing machineries^1,3,5–7^, providing a distinct biochemical environment that either enhances reaction rates or suppresses processing^8,9^. Consequently, the functional impact of an RNP condensate on an RNA molecule is influenced by the RNA’s residence time before escaping the condensate. Despite its importance, the mechanisms regulating RNA residence time remain poorly understood, partly due to the lack of knowledge about internal architectures within condensates, where locally tuned diffusion could influence RNA retention and processing.

To address this challenge, we developed a multi-tether single-molecule tracking (SMT; used here synonymously to single-particle tracking, SPT) strategy that enables precise, simultaneous tracking of fluorophore-labeled RNA and protein molecules within reconstituted, full-length, tag-free Fused in Sarcoma (FUS) condensates. We selected FUS as a model system because, as a protein with extensive intrinsically disordered regions (IDRs), it forms a variety of physiological and pathological condensates^10–14^. Using this approach, we discovered that a significant fraction of RNA and protein molecules are confined within distinct nanometer-scale domains, which are sub-condensate architectures that restrict diffusion and extend residence time. These nanodomains dynamically regulate molecular interactions, with the potential to shape RNA processing and protein activity within condensates.

Furthermore, we found that nanodomains influence the liquid-to-solid phase transition of FUS condensates, a process known as condensate aging, which results in FUS fibrilization inside neurons and is implicated in amyotrophic lateral sclerosis (ALS) and other neurodegenerative diseases^13,15–19^. During in vitro condensate aging, we observed that FUS nanodomains migrate toward the condensate periphery, providing a favorable chemical environment for FUS fibrilization, consistent with prior reports of fibers forming at condensate surfaces^13,20^. We also discover that the two FDA-approved ALS drugs edaravone and riluzole render nanodomains more susceptible to these aging-induced changes. Our findings establish a direct link between nanoscale condensate sub-structures and pathological aggregation, highlighting nanodomains as key regulators of condensate maturation and disease-related transitions.

### Single molecule tracking reveals nanodomains in full-length FUS condensates

Largely disordered RNA-binding proteins, such as FUS, play key roles in nuclear RNA processing and transport while also acting as scaffold molecules that drive phase separation, with RNAs partitioning into the resulting condensates as guest molecules^3,6,7^. While phase separation studies often focus on the N-terminal prion-like domain (PrLD, 1–239 aa) of FUS^21–24^, this approach neglects the structural and electrostatic contributions of the C-terminal regions (Fig. 1a,b), which may significantly influence condensate properties. To better replicate physiological conditions, we purified full-length, purification tag-free FUS as described^25,26^ (Fig. S1) and synthesized a ∼1,500 nucleotide (nt) firefly luciferase (FL) mRNA containing features essential for intracellular translation (Fig. 1a and Supplementary Fig. 1). For fluorescence-based SMT, we employed click-chemistry to covalently attach minimally disruptive^27^ organic dye molecules between the coding sequence and unmodified poly(A) tail (Fig. 1a and Supplementary Fig. 1), whereas sparsely labeled FUS via surface lysines, and doped low concentrations of both biomolecules into condensates of unlabeled FUS.

**Fig. 1.**
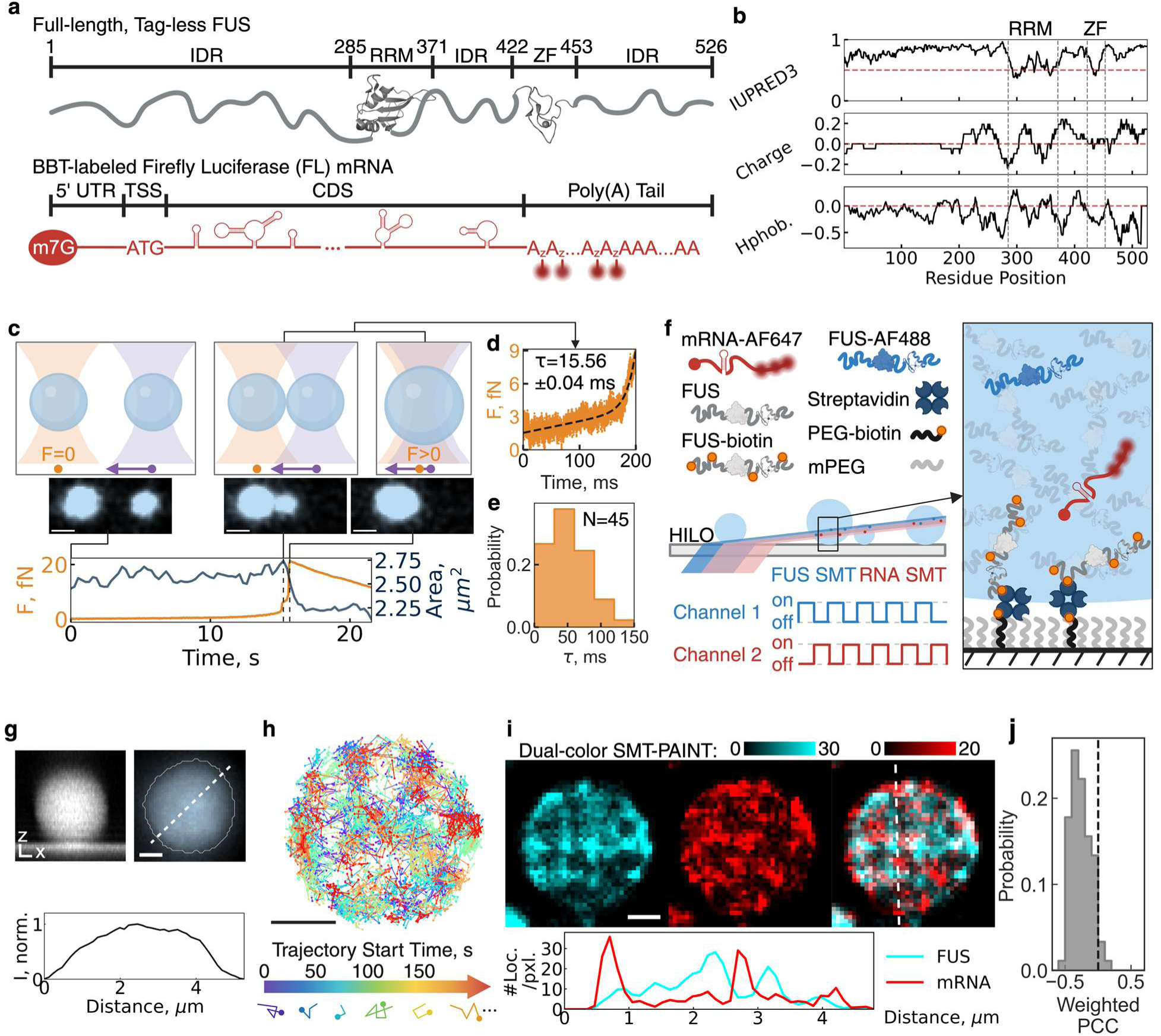
Tethering enables accurate single particle tracking of protein and RNA molecules within 3D biomolecular condensates. **a** Schematic of the purified model protein and RNA. **b** Physicochemical profile of the full-length FUS protein. All curves are running window analysis with a window size of 21 residues. IUPRED3 is a prediction score for IDR, curated by experimental database DisProt. Charge density is calculated from each residue’s pKa assuming pH=7. Hydropathy (Hphob.) is calculated with hydrophobicity scales determined by Eisenburg et al^57^. The red dotted line indicates the threshold of structured (<0.5) versus unstructured (>0.5), positively versus negatively charged, or hydrophobic (>0) versus hydrophilic (<0). **c** Schematic of a C-Trap experiment that combines the force measurement from optical tweezers and the condensate total area from scanning-confocal microscope to measure the rheology of full-length, tag-free FUS condensates. **d** Fitting high-frequency force data to determine the rate constant *τ* (characteristic fusion time) of condensates. **e** Distribution of *τ*. Measurements were done in three biological replicates with N being the number of condensate pairs measured. **f** Schematic of a tethered, sparsely labeled condensate system enabling single-molecule tracking (SMT) in one or two channels simultaneously within a single biomolecular condensate. #1-3 are three types of experiments to determine the sub-condensate heterogeneity with different probes. For bulk fluorescence imaging of condensates, a labeling ratio < 0.01% was used to minimize the impact of fluorophores on condensates. **g** Side view of a representative tethered FUS condensate shows its spherical shape in 3D. Bulk fluorescence image of tethered FUS condensates under highly inclined laminated optical light sheet (HILO) microscope with condensate boundary detected by machine-learning-based pixel classifier, the best method for condensates with a broad range of sizes^58^, and the normalized intensity (I, norm.) profile along the white dotted line of the zoomed-in condensate. **h** Raw SMT trajectories of FUS molecules within a single condensate and schematic for two types of trajectory reconstruction. A plot of all FUS trajectories in a representative tethered condensate, colored by the start time of each trajectory. A trajectory localization density map is formed by accumulating locations of all trajectories throughout a SMT dataset, while a time-lapse trajectory reconstruction video is reconstructed by accumulating locations of trajectories within a non-overlapping time window into frames of a video. **i** Representative RNA trajectories within a condensate. Dual-color trajectory reconstruction of FUS and mRNA with a cross-sectioning profile across the white dotted line. Pixel intensity represents the number of SMT trajectories on each pixel. **j** Distribution of pixel-wise Pearson Correlation Coefficient (PCC) of **e**, weighted by the number of SMT trajectories per pixel. A gray dotted line indicates PCC=0, distinguishing positive correlation (PCC>0) and negative correlation (PCC<0). IDR, intrinsically disordered region; RRM, RNA-recognition motif; ZF, zinc-finger domain; BBT, between-body-and-tail labeling strategy for mRNA; UTR, untranslated region; TSS, translation start site; CDS, coding sequence; ALEX, alternating-laser excitation; PEG, polyethylene glycol; mPEG, PEG monomethyl ether. All scale bars are 1 µm.

In this model system, we induced FUS condensation (and Supplementary Fig. 2) at a near-physiological concentration of 10 µM^13^. Using microrheology in a dual-trap optical tweezers setup (Fig. 1c), we first measured the condensate fusion kinetics, extracting the characteristic fusion time *τ* from the force curve as droplets transitioned from initial contact to full fusion^13,28^ (Fig. 1d). Notably, full-length tag-free FUS exhibited significantly faster fusion (*τ*_*mean*_ = 51 ± 4 ms) (Fig. 1e) compared to tagged variants, including green fluorescence protein-appended FUS (*τ*_*mean*_ = ∼150 ms)^13^, and maltose-binding protein-fused FUS (*τ*_*mean*_ = ∼570 ms)^28^. These results indicate that full-length, tag-free FUS forms highly dynamic, liquid-like condensates with distinct biophysical properties.

Next, we introduced 10 nM sparsely Alexa Fluor 488-labeled FUS during condensation and utilized highly inclined and laminated optical sheet (HILO) microscopy to visualize the entire condensate (Fig. 1f). We devised a surface multi-tethering approach that enables intra-condensate single-molecule fluorescence tracking of both RNA and protein molecules within liquid-like condensates, providing a foundation for studying sub-condensate organization (see Methods). The surface-captured FUS condensates maintained a spherical shape in all dimensions (Fig. 1g and Supplementary Fig. 3), in contrast to the previously reported wetted flat domains^29,30^. Furthermore, the uniform fluorescence intensity within tethered condensates (Fig. 1g) confirms that FUS forms a single-phase condensate, unlike the multiphase RNP condensates observed in prior studies^31^. These findings establish this system as an ideal in vitro mimic of physiological FUS condensates for high-precision SMT.

For single-molecule tracking (SMT), we incorporated 50 pM fluorescently labeled FUS protein and/or FL mRNA into the condensation reaction and performed dual-color HILO microscopy with alternating laser excitation (ALEX) to suppress channel cross-talk (Fig. 1f). To optimize trajectory length, we included an oxygen scavenging system (OSS), leading to a concentration-dependent increase in trackable particles (and Supplementary Fig. 4). This allowed us to generate cumulative single-protein and RNA trajectory maps within individual condensates (Fig. 1h).

In these maps, fast-moving molecules produce fewer localizations and thus less dense point accumulations, whereas confined molecules generate higher local point densities (Fig. 1i). By reconstructing localization density maps from freshly prepared FUS condensates, we identified fast-diffusing regions as low-intensity signals and slow-diffusing regions as high-intensity signals, revealing nanometer-scale regions—termed nanodomains—that restrict both FUS and mRNA diffusion (Fig. 1i and Supplementary Fig. 5). Notably, FUS and mRNA nanodomains exhibited a significant mutual exclusion effect, as reflected by a negative mean pixel-wise Pearson correlation coefficient (PCC) of −0.26 ± 0.02 (Fig. 1j), confirming that they are structurally distinct. Since no protein density transition was observed (Fig. 1g), these nanodomains likely represent diffusion-limiting percolation networks rather than phase-separated subdomains. Our findings align with previous observations reporting non-uniform localization densities within biomolecular condensates using small-molecule dyes such as Nile Red^32^. (Of note, in our experiments, Nile Red aggregated when partitioning into FUS condensates (Supplementary Fig. 6)).

### RNA molecules are retained in nanodomains dependent on length and charge

To systematically characterize heterogeneous intra-condensate diffusion behaviors (Fig. 2a), we developed a computational pipeline to classify SMT trajectories into immobile, confined diffusion, or normal (Brownian) diffusion states (Fig. 2a-b and Supplementary Fig. 7). We distinguished initial categories using the mean step size of a trajectory and the anomalous diffusion component α^33^ calculated from mean squared displacement (MSD)-lag time (*τ*) (Fig. 2a-b). To assess population-wide shifts, we applied a Bayesian statistics-based state array (SA) method^34^ (Fig. 2c) to estimate the distribution of apparent diffusion coefficient (D_app_). The diffusion profile of FL mRNA in the dilute phase closely matched that of a random walk simulation (Supplementary Fig. 8) and of 20-nm beads (Fig. 2c and Supplementary Fig. 9), consistent with the 20-nm radius of gyration previously determined for a 1,500-nt RNA^35^. This agreement validates our pipeline’s accuracy for the purpose of intra-condensate RNA diffusion dynamics and provides a threshold for classifying molecules as confined or normal diffusion based on the anomalous diffusion component α.

**Fig. 2.**
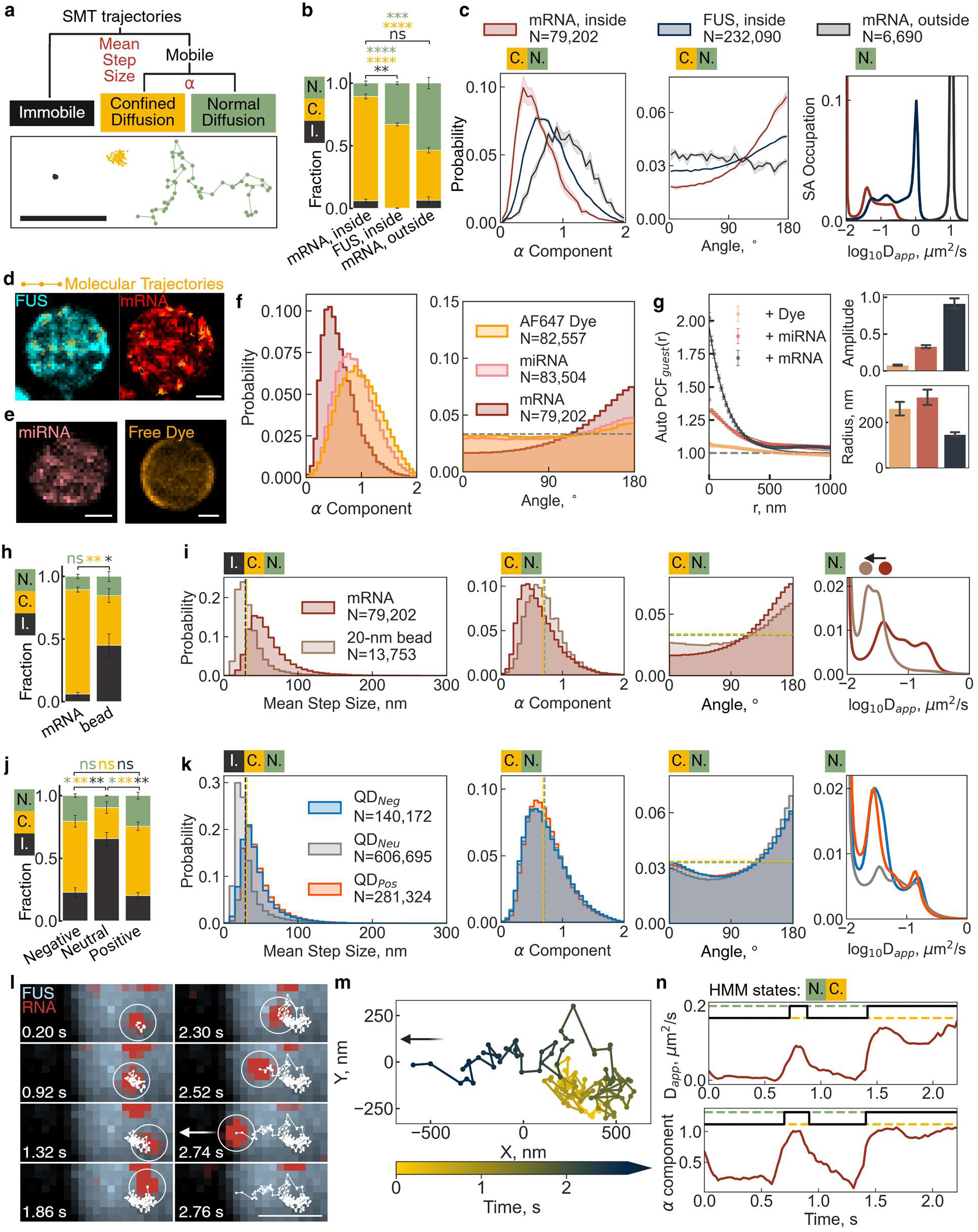
FUS, miRNA, and mRNA are confined in different nanodomains based on physicochemical properties. **a** Three representative mRNA trajectories within a single full-length tag-free FUS condensate, which can be classified into three categories, immobile, confined diffusion, and normal diffusion. **b** Fractions of three categories for mRNA and FUS molecules in the condensed phase and mRNA molecules in the dilute phase. Error bars are S.E.M. from more than three experiments on different dates. **c |** Distributions of the anomalous diffusion component α, the angle between adjacent steps, and the apparent diffusion coefficient (D_app_). The α and angle distributions are from all mobile molecules while the D_app_ distribution is from only normal diffusion molecules. Transparent color represents the S.E.M. from more than three experiments on different dates. D_app_ distribution is calculated by a state array (SA) method^60^, while the D_app_ distribution calculated by mean squared displacement (MSD)-*τ* fitting could be found in Fig. S7. D_app_ distribution is also truncated to visualize the smaller peaks. So as in **i** and **k**. **d** Localization density maps of trajectory locations, FUS in cyan and mRNA in red, with representative trajectories overlaid in orange. **e** Localization density maps of trajectory locations of miRNA in red and Alexa Fluor 647 in gray. **f** Distributions of the anomalous diffusion component α and the angle between adjacent steps. **g** Plot of calculated autocorrelation (g(r)^guest^) of guest molecules represented in **d** and **e**. **h, j** Fractions of the immobile (I.), confined diffusion (C.) and normal diffusion (N.) categories for mRNA, beads, and quantum dots in the condensed phase. (**h**), mRNA molecules and 20-nm carboxylate-modified beads in the condensed phase (**j**), and amine-, PEG-, carboxylate-functionalized quantum dots (QDs) in the condensed phase. Error bars are S.E.M. from more than three experiments on different dates. **i, k** Distributions of the mean step size of each trajectory, the anomalous diffusion component α, the angle between adjacent steps, and the D_app_ values for mRNA, miRNA, and free Alexa Flour 647 dye molecules in the condensed phase (**f**), mRNA molecules and 20-nm carboxylate-modified beads in the condensed phase (**i**), and amine-, PEG-, carboxylate-functionalized quantum dots (QDs) in the condensed phase (**k**). The α and angle distributions are from all mobile molecules or particles while the D_app_ distribution is from only normal diffusion molecules or particles. D_app_ distribution is calculated by a state array (SA) method, while the D_app_ distribution calculated by mean squared displacement (MSD)-*τ* fitting could be found in Fig. S7. D_app_ distribution is also truncated to visualize the smaller peaks. A double-layered black-yellow dotted line indicates the threshold between immobile (black) and confined diffusion (yellow) diffusion, while a double-layered yellow-green dotted line indicates the the threshold between confined (yellow) and normal (green) diffusion. **l** Example multi-stage diffusion trajectory of an mRNA moving from one confined region to another, following by normal diffusion until it leaves the tethered FUS condensate from the edge. The white circle indicates the current position of mRNA while the white line indicates the trajectory before the current frame. **m** A zoomed-in view of the trajectory, colored by the time each location is detected. **n |** Running-window analysis of D_app_ and α value from a window size of 20 steps. Hidden Markov Model (HMM) is used to determine the transition point between confined and normal diffusion states. All scale bars are 1 µm. Statistics annotation: independent t-test, ns: 0.05 < p <= 1, *: 0.01 < p <= 0.05, **: 0.001 < p <= 0.01, ***: 0.0001 < p <= 0.001, ****: p <= 0.0001.

Applying our SMT analysis pipeline to FUS and RNA diffusion within condensates revealed multiple normal diffusive states of FUS with the major peak of the D distribution located at 0.8 μm²/s (Fig. 2c and Supplementary Table 1). Previous studies of membrane-wetted domains and non-tethered 3D condensates similarly reported multiple normal diffusion states of scaffold proteins, but with low D_app_ values ranging from ∼0.01 to ∼0.52 μm²/s^29,36–38^.

We found that the movement of both FUS and mRNA molecules was highly restricted, with 67 ± 1% of FUS and 83 ± 2% of mRNA exhibiting confined diffusion, compared to only 40 ± 2% for mRNA in the dilute phase (Fig. 2b) and 32% for simulated random walk trajectories (Supplementary Fig. 8). FUS and mRNA confinement was further supported by α component distributions skewing toward 0 and angle distributions trending toward 180°, yielding mean α and angle values of 0.81 ± 0.01 and 104.6 ± 0.4° for FUS, and 0.61 ± 0.02 and 117.6 ± 1.5° for mRNA, respectively (Fig. 2c and Supplementary Table S1). For non-confined FUS and mRNA molecules, their partially overlapping D_app_ distributions are consistent with the formation of RNP complexes, between the sparse mRNA guest molecules and the large excess of FUS, whereas the fast-diffusing peak unique to FUS reflects the smaller molecular weight of free FUS (Fig. 2c). Additionally, the major D_app_ peak of normal diffusive mRNA within condensates was slower than the 10 μm²/s D_app_ peak of mRNA in the dilute phase, in agreement with the expected molecular crowding in the condensed phase.

Overlaying single-molecule trajectories classified as confined diffusion with our localization density maps further confirmed that nanodomains serve as regions of restricted molecular diffusion (Fig. 2d). In contrast, Cy5-labeled 21-nt microRNA miR-21 and free Alexa Fluor 647 dye exhibited progressively less confinement than FL mRNA, with the free fluorophore displaying primarily normal diffusion, aside from some confinement near the condensate surface (Fig. 2e,f). Using a pair correlation function, we found that the nanodomain clustering amplitude and size (radius) were RNA length-dependent, with mRNA-containing nanodomains appearing smaller and more tightly clustered than those formed by both miRNA and dye molecules (Fig. 2g).

To further investigate the mechanisms governing nanodomain size and stability, we compared FL mRNA diffusion with carboxylate-modified 20-nm beads inside FUS condensates (Fig. 2h-i). The spherical beads exhibited a higher immobile fraction, consistent with their inability to thread through a meshwork-like environment, such as a percolated condensate^39^. In contrast, the mRNA’s ability to thread through such a condensate meshwork likely explains its faster normal diffusion compared to the beads (Fig. 2i).

To assess the role of surface charge, we examined 9.5-nm quantum dots (QDs) with carboxylate (negative), PEG (neutral), or amine (positive) surface functionalization (Fig. 2j-k). Surprisingly, surface charge had no significant effect on the diffusion profiles (Fig. 2j), but neutral QDs exhibited a significantly larger immobile fraction than their charged counterparts (Fig. 2k). These findings support the notion that a RNA’s size determines its confinement within nanodomains, while both a non-spherical shape and negative charge facilitate mobility within condensates.

We next investigated whether confinement affects the intra-condensate residence time of selectively trapped RNA molecules. Indeed, we found that the escape trajectory of a single mRNA molecule was prolonged by transient nanodomain retention (Fig. 2l). A representative trajectory revealed that the mRNA molecule was confined within two separate nanodomains— first from 0 to 0.6 s, then from 0.9 to 1.4 s—before reaching the condensate boundary at 2.74 s (Fig. 2m), where rapid axial diffusion in the dilute phase led to signal loss. A running-window analysis of D_app_ and α confirmed transitions between normal and confined diffusion states (Fig. 2n). This extension of residence time by association with nanodomains suggests a potential regulatory role for nanodomains in modulating mRNA processing efficiency both inside and outside of protein condensates by selective sequestration.

### RNA guest molecules modulate FUS nanodomains

Next, we investigated whether, conversely, RNA guest molecules influence the properties of scaffold protein nanodomains. Analyzing localization density maps of FUS molecules within condensates in the presence of mRNA, miRNA (Supplementary Fig. 10), or dye guest molecules revealed that guest molecules generally weaken FUS nanodomains (Fig. 3a). Guest-free FUS condensates exhibited the strongest nanodomain clustering amplitude (1.0 ± 0.2) and the largest nanodomain radius (289 ± 59 nm), both of which diminished to varying degrees upon the addition of guest molecules (Fig. 3a). Additionally, we observed that as guest molecule size increased, the normal diffusion population of FUS shifted toward slower diffusion (Fig. 3b and Supplementary Fig. 11). Radial distribution analysis of FUS nanodomains in the presence of guest molecules within condensates revealed a significantly higher probability of nanodomain localization at the condensate surface (Fig. 3c), suggesting a global structural influence of guest molecules on condensate organization. In contrast, radial distributions of guest molecules showed dye enrichment at the condensate interface, while RNA molecules remained uniformly distributed (Fig. 3d).

**Fig. 3.**
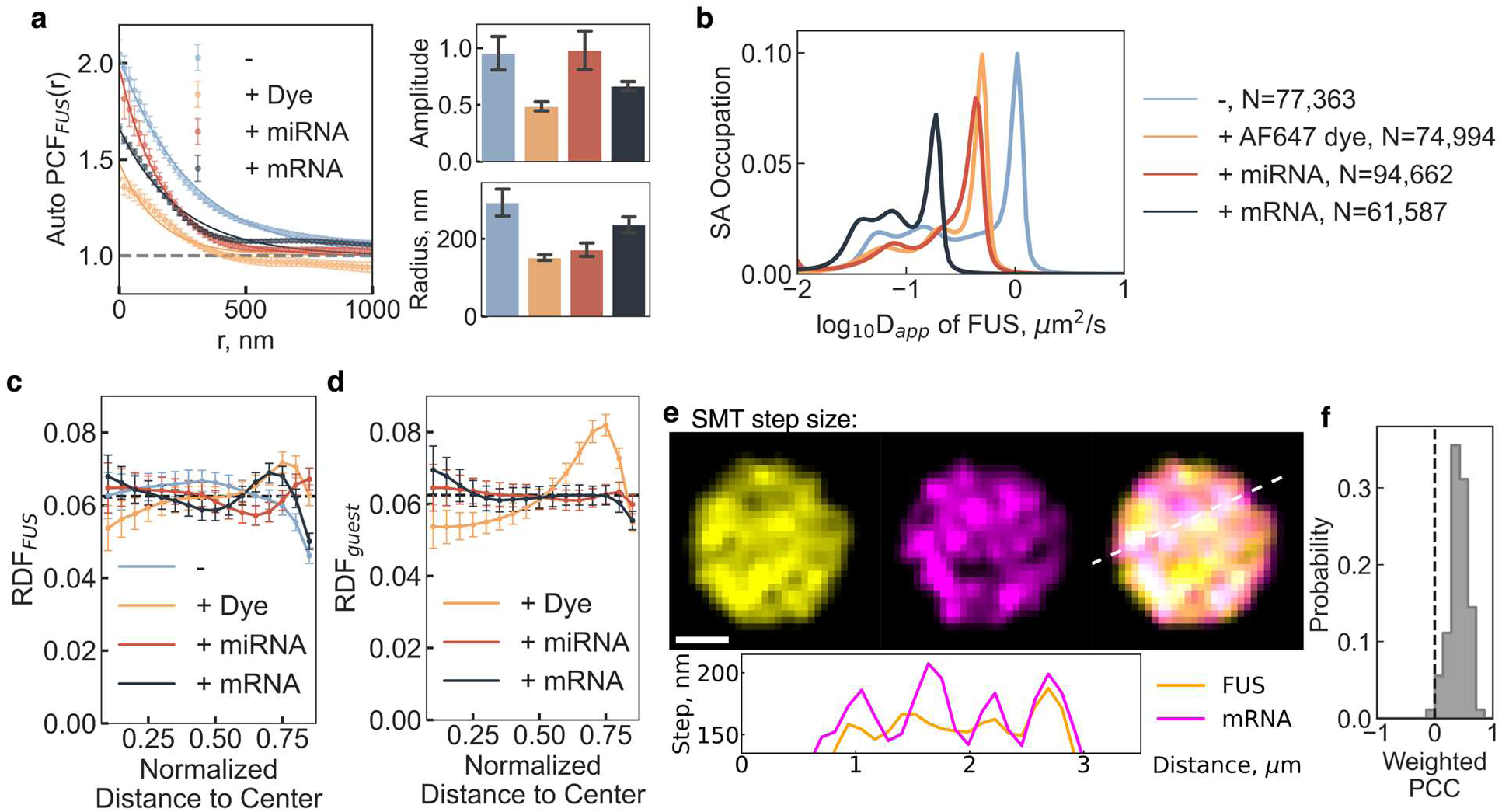
FUS nanodomains formation is tuned by partitioned guest molecules. **a** Plot of calculated autocorrelation (g(r)^FUS^) of scaffold molecules, FUS**. b**D_app_ distribution of FUS calculated by the SA method. **c** Plot of radial distribution of FUS nanodomains within condensates of varying guest molecules. **d** Plot of radial distribution of guest nanodomains within condensates. **e** Average step size on each pixel for FUS and mRNA trajectories with a cross-sectioning profile across the white dotted line. higher the pixel intensity, higher the average step size, faster the diffusion. **f** Pixel-wise weighted PCC of **e**. Statistics annotation: independent t-test, ns: 0.05 < p <= 1, *: 0.01 < p <= 0.05, **: 0.001 < p <= 0.01, ***: 0.0001 < p <= 0.001, ****: p <= 0.0001.

To examine the spatial distribution of unconfined molecules, we analyzed step size heatmaps to map fast-diffusing molecules within condensates (Fig. 3e). We found that fast-diffusing FUS and mRNA molecules were spatially correlated (Supplementary Fig. 12), with a positive mean PCC of 0.42 ± 0.02 (Fig. 3f) and a positive cross-pair correlation (G_cross_; Supplementary Fig. 13). This observation suggests fast diffusion of both protein and RNA molecules outside of nanodomains, independent of the formation of RNP complexes.

Collectively, these findings support a model of intra-condensate diffusion in which nanodomains selectively trap scaffold proteins and RNA guest molecules (Fig. 1i-j), while the same biomolecules, when fast-diffusing and unconfined, remain excluded from these trapping regions (Fig. 3e-f and Supplementary Fig. 13). Intriguingly, RNA guest molecules not only weaken FUS nanodomains but also promote their localization to the condensate periphery, further influencing condensate architecture and dynamics.

To further explore the molecular basis of nanodomain organization, we analyzed FUS using CIDER^40^, a well-established framework for characterizing intrinsically disordered regions (IDRs) in protein sequences. This analysis revealed that the N-terminal IDR (aa1-285) displays a segregated charge distribution favoring globular conformations, whereas the C-terminal IDR (aa453-526) exhibits a more uniform charge distribution, consistent with a broader ensemble of structural states, including both expanded and collapsed conformations (Supplementary Fig. 14). We hypothesize that this conformational heterogeneity underlies the emergence of two distinct classes of nanodomains: high-connectivity regions that confine expanded FUS molecules, and low-connectivity regions in which large, extended RNA molecules scaffold collapsed FUS molecules into a “beads-on-a-string” arrangement.

Supporting this model, prior work has shown that RNAs, such as FL mRNA, can adopt both expanded polymer conformations and compact states with short end-to-end distances^41^, enabling them to act as scaffolds for percolated FUS networks. The high multivalency of expanded RNAs enhances their ability to engage the FUS meshwork, facilitating stable confinement within RNA-rich nanodomains. Together, our findings point to a highly organized intra-condensate landscape, in which proteins and RNAs are selectively confined to distinct nanodomains, while faster-diffusing, likely more globular species remain excluded from these regions.

### Nanodomains facilitate FUS surface aggregation through liquid-to-solid transition

Intrigued by the peripheral localization of FUS nanodomains induced by guest molecules (Fig. 3c) and our observation of surface-originating fibril outgrowth from FUS condensates after 24 hours (Fig. 4a), we hypothesized that confined diffusion near the condensate surface may be linked to condensate aging^13,42^. Applying our trajectory localization density map and diffusion analysis pipelines to aged condensates, we observed a higher incidence of surface-localized nanodomains at 8 and 24 hours of aging (Fig. 4b), consistent with the idea that FUS nanodomains facilitate the liquid-to-solid transition of FUS into Fibrils^20,42,43^.

**Fig. 4.**
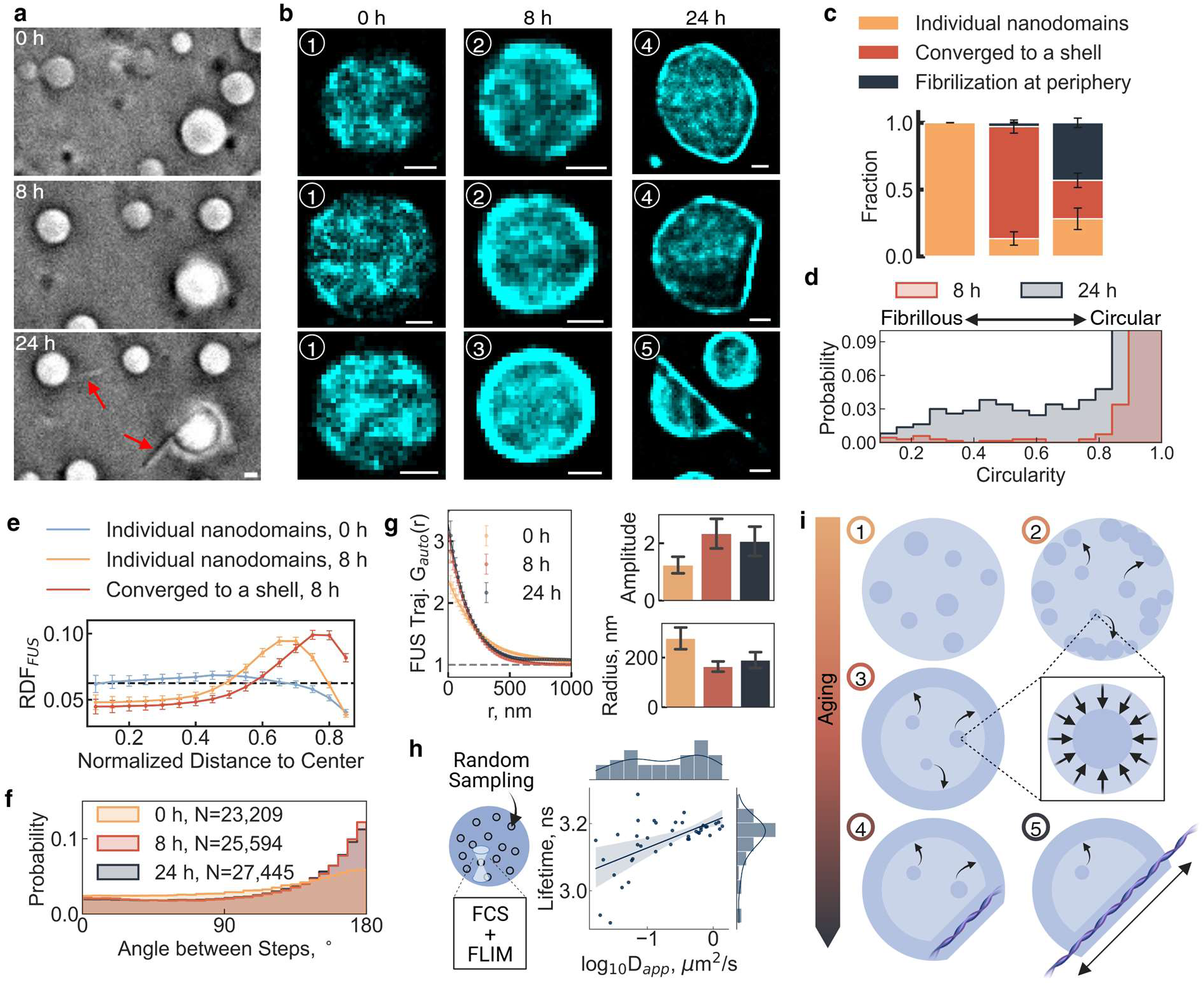
Nanodomains colocalize with FUS fibrilization sites. **a** Bright-field images of FUS fibril out-growth from condensate surface after 24 hours incubation at room temperature. **b** Representative localization density maps of condensates at 0-, 8-, and 24-hour time points. Numbers 1-5 represent morphological classification of condensates in each group related to the stage of phase transition from liquid-to-solid. **c** Bar plot of morphological changes that emerge at 0-, 8-, and 24-hour time points. **d** Plot of circularity changes of condensates from 8- to 24-hour time points. **e** Plot of radial distribution of FUS nanodomains within condensates at 0-, 8-, and 24-hour time points. **f** Plot of angle between steps for trajectories within condensates at 0-, 8-, and 24-hour time points. **g** Plot of calculated autocorrelation (g(r)^FUS^) of FUS molecule trajectories at 0- and 8-hour time points represented in **b**. **h** Cartoon and results of a random-sampling fluorescence correlation spectroscopy (FCS)-FLIM assay. The scatter plot shows the correlation between D_app_ measured by FCS and lifetime by FLIM with the distributions of each on each side. The blue line shows a linear regression model fit with the shaded area being the 95% confidence interval of the fit. **i** Proposed model of the role of nanodomains in aging and the liquid-to-solid transition of FUS. (1) Nanodomains uniformly distributed, (2) most nanodomains move to periphery while interior nanodomain clustering diminishes, (3) nanodomains peripherally accumulate into shell, (4) fibrilization seeded, (5) fibril outgrowth. All scale bars are 1 µm. Statistics annotation: independent t-test, ns: 0.05 < p <= 1, *: 0.01 < p <= 0.05, **: 0.001 < p <= 0.01, ***: 0.0001 < p <= 0.001, ****: p <= 0.0001.

In our localization density maps, we identified three major morphological classes: individual nanodomains, nanodomains peripherally converged to a shell, and fibrilization at the condensate surface (Fig. 4b,c). At 0 hours, condensates contained only individual nanodomains. By 8 hours, most condensates exhibited nanodomain convergence to the periphery, while by 24 hours, immobile fibrilization became the predominant morphology, accompanied by a significant loss of condensate sphericity (Fig. 4c,d). Radial distribution analysis confirmed these classifications, demonstrating a strong preference for nanodomain localization at the periphery after aging (Fig. 4e). Additionally, angle distribution analysis revealed an increase in molecular confinement at 8 hours (Fig. 4f). Further, both nanodomain radius and amplitude trended toward smaller and tighter clusters at 8 and 24 hours, reinforcing their role in driving condensate maturation and fibrilization (Fig. 4g).

We hypothesized that nanodomains facilitate the liquid-to-solid transition by creating a unique chemical environment that modulates local diffusion and seeds fibril formation. To test this notion, we used fluorescence lifetime imaging (FLIM) combined with fluorescence correlation spectroscopy (FCS) to measure D_app_ at ten random locations within individual condensates, revealing a positive correlation between fluorescence lifetime and D_app_ (Fig. 4h). Additionally, we observed a ∼0.3 ns decrease in fluorescence lifetime via FLIM (Supplementary Fig. 15).

As a control, we found that mRNA confinement was only marginally affected by condensate aging (Supplementary Fig. 16). Similarly, drastically increasing the RNA concentration by adding 50 ng/μL unlabeled total HeLa cell RNA during condensate reconstitution had little effect on freely diffusive mRNA and did not alter its nanodomain-induced confinement (Supplementary Fig. 17).

Our findings further reveal that FUS nanodomains exhibit slow mobility and, as condensates age, become increasingly localized at the periphery (Supplementary Fig. 15), where a distinct chemical environment promotes local aggregation and fibrilization (Supplementary Fig. 18).

Strikingly, the factors governing molecular mobility and confinement within nanodomains are distinct. Globular yet irregularly shaped molecules with high charge density, such as long mRNAs, but less so short miRNAs, can thread through the condensate network, remaining mobile; however, upon further expansion, multivalent interactions begin to anchor them to nanodomains. In contrast, globular particles like beads exhibit reduced mobility due to steric hindrance, particularly when charge-neutral. By comparison, the mixed distribution of charged residues in full-length FUS (Fig. 1b) may act as a “molecular grease,” promoting transient interactions and facilitating mobility, which becomes restricted in nanodomains only upon structural expansion. This framework helps explain why previous studies, which focused on truncated or tagged IDR proteins, may have overlooked nanodomain-confined diffusion.

Together, these findings indicate that the chemical environment within aging FUS nanodomains specifically slows protein diffusion, thereby promoting local aggregation and fibrilization, while having minimal impact on RNA mobility (Fig. 4i). Our findings nevertheless align with previous reports of condensate heterogeneity and the coupling of percolation with phase separation^1^. They additionally reveal, however, that percolation occurs locally rather than globally, emerging immediately after condensate assembly to influence sub-condensate organization and molecular residence time (Fig. 4b). While intra-condensate heterogeneity driven by multiphase architectures, such as in the nucleolus, has been extensively studied, examples of heterogeneity independent of secondary phase transitions have only recently been reported^29,42^. Recent bulk fluorescence analyses of condensed FUS molecules provide evidence of intra-condensate heterogeneity similar to our FLIM analysis^44^. Among the various physical transitions governing phase separation, percolation is one of the best-studied, playing key roles both in the pre-condensation assembly of protein clusters below the saturation concentration and in the organization of protein-RNA condensates after their formation^1^.

### Small molecule drugs enhance nanodomains and condensate aging

Given that protein nanodomains regulate the surface-induced liquid-to-solid transition and fibrilization of FUS, and that in turn even small guest molecules were found to modulate FUS nanodomains (Fig. 3c), we investigated the impact of two FDA-approved small-molecule drugs for ALS, a neurodegenerative disease characterized by pathological cytoplasmic FUS fibrilization in motor neurons^19,45,46^. When FUS condensates were prepared in the presence of micromolar concentrations of edaravone or riluzole—both previously shown to partition into biomolecular condensates^47^—nanodomains became more susceptible to aging-induced relocation toward the condensate surface (Fig. 5a). Notably, confined FUS nanodomains most consistently shifted to the surface in the presence of riluzole, while diffusion state analysis revealed that unconfined FUS molecules still exhibited fast diffusion (Fig. 5c). Supporting the idea that surface-localized nanodomains promote fibrilization, bright-field imaging of 24-hour-old condensates showed that riluzole enhanced fibril formation from the condensate surface (Fig. 5b and Supplementary Fig. 19).

**Fig. 5.**
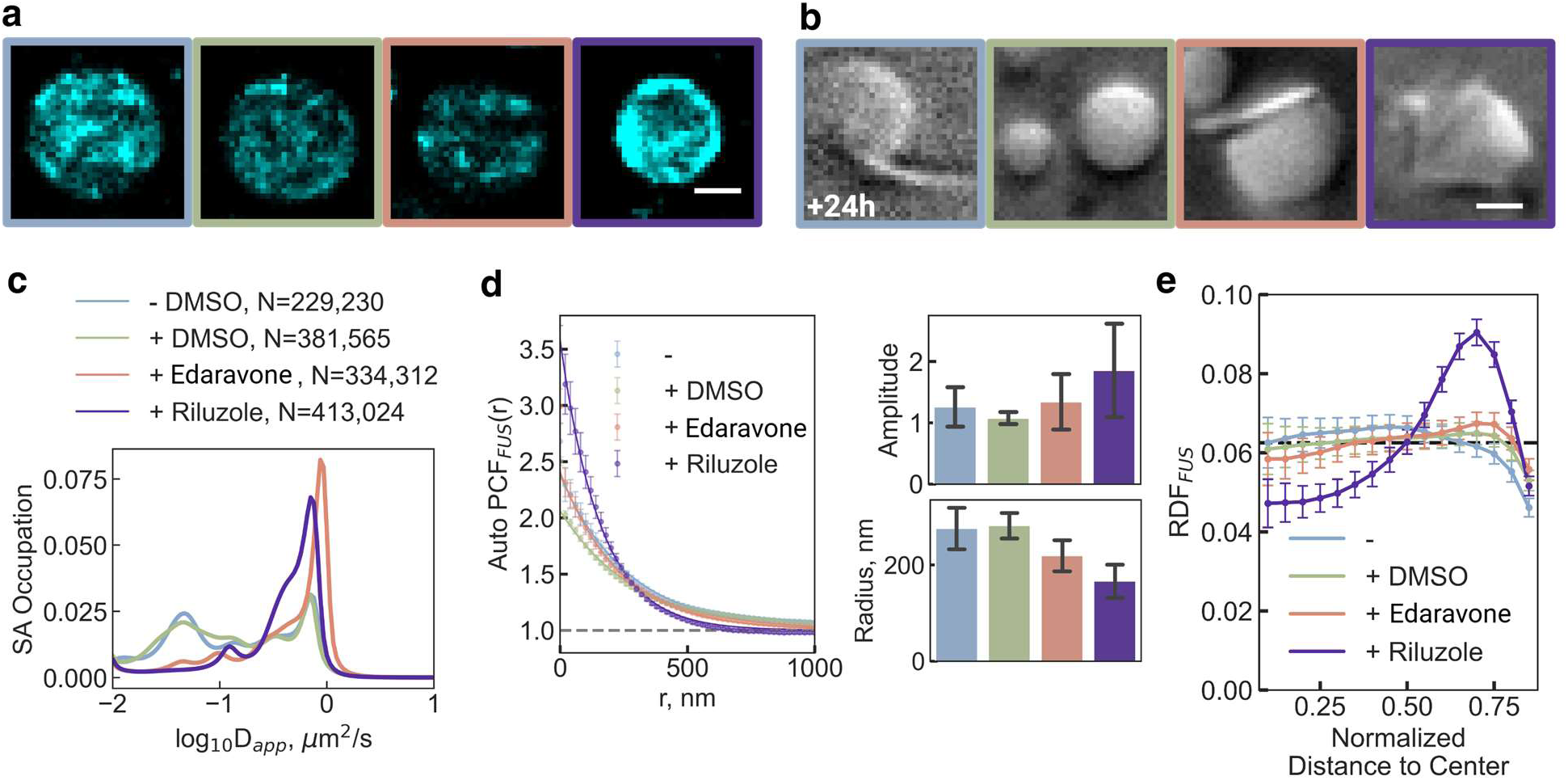
Small molecule drugs can affect FUS nanodomain formation. a Representative localization density maps of condensates in the presence of either edaravone or riluzole ALS drug. Border colors represent corresponding small molecule added to the condensate: blue, no addition; green, DMSO; salmon, edaravone; purple, riluzole. b Representative bright field images of 24-hours-aged condensates under drug inclusion. c D_app_ distribution of FUS calculated by the SA method. d Plot of calculated autocorrelation (g(r)^FUS^) of FUS molecule trajectories under various drug treatments. Corresponding bar plots compare amplitude and radius of the nanodomains. e Plot of radial distribution of FUS nanodomains within condensates under various drug treatments.

Auto-PCF analysis confirmed that, compared to a DMSO solvent control, both edaravone and riluzole maintained or increased the nanodomain correlation amplitude while reducing the radius (Fig. 5d), mirroring effects observed with other guest molecules such as fluorescent dyes (Fig. 3a). Radial distribution analysis further revealed that the peripheral localization of nanodomains was only weakly enhanced by edaravone yet significantly promoted by riluzole (Fig. 5e). These nanodomain rearrangements directly influence molecular diffusion, likely creating localized environments amenable to fibril formation (Fig. 5b).

The observation that edaravone and riluzole accelerate aging-induced nanodomain remodeling suggests a potential connection between these small-molecule drugs and the liquid-to-solid transition of FUS, beyond their established roles as a free radical scavenger and inhibitor of glutamate excitotoxicity, respectively^45–47^. Although riluzole is primarily known to reduce presynaptic glutamate release and dampen neuronal excitability^46^, our findings align with emerging evidence work linking excitotoxic stress and FUS-related pathology^48^ and demonstrate that small aromatic compounds can modulate protein aggregation^49^. Together, these insights offer a more nuanced understanding of riluzole’s molecular mechanisms and highlight the importance of continued investigation. More broadly, our results emphasize that nanodomain dynamics play a critical role in condensate behavior, providing new perspectives on how small molecules might modulate the pathological properties of biomolecular condensates.

## Conclusions

Nanodomains independently slow the diffusion of protein and RNA molecules, thereby influencing their biochemical reaction capacity (Fig. 6). Unlike secondary phase separation, nanodomain formation arises from local percolation among biomolecules and―in the case of full-length FUS―is modulated by guest molecules. Our findings suggest that nanodomains play a key role in regulating molecular residence time and in facilitating the liquid-to-solid transition and fibrilization of FUS (Fig. 6), a process linked to neurodegenerative diseases such as ALS. The high connectivity within FUS nanodomains likely facilitates inter-chain interactions, potentially priming the surface-originating fibril growth of condensates that we observe. Our observations are consistent with recent studies showing that protein molecules retain fast structural dynamics^50^ within condensates, suggesting that fibrilization then arises from enhanced intermolecular connectivity through percolated nanodomains rather than from changes in intramolecular dynamics and global percolation. Notably, the FDA-approved ALS drugs edaravone and riluzole both accelerate aging-related FUS—yet not RNA—nanodomain relocalization to the condensate shell, where the protein becomes prone to fibrilization. These findings suggest that small-molecule drugs may play previously unrecognized roles in ALS pathogenesis, potentially protecting neurons by promoting fibril formation as a sink for toxic intermediates^51^.

**Fig. 6.**
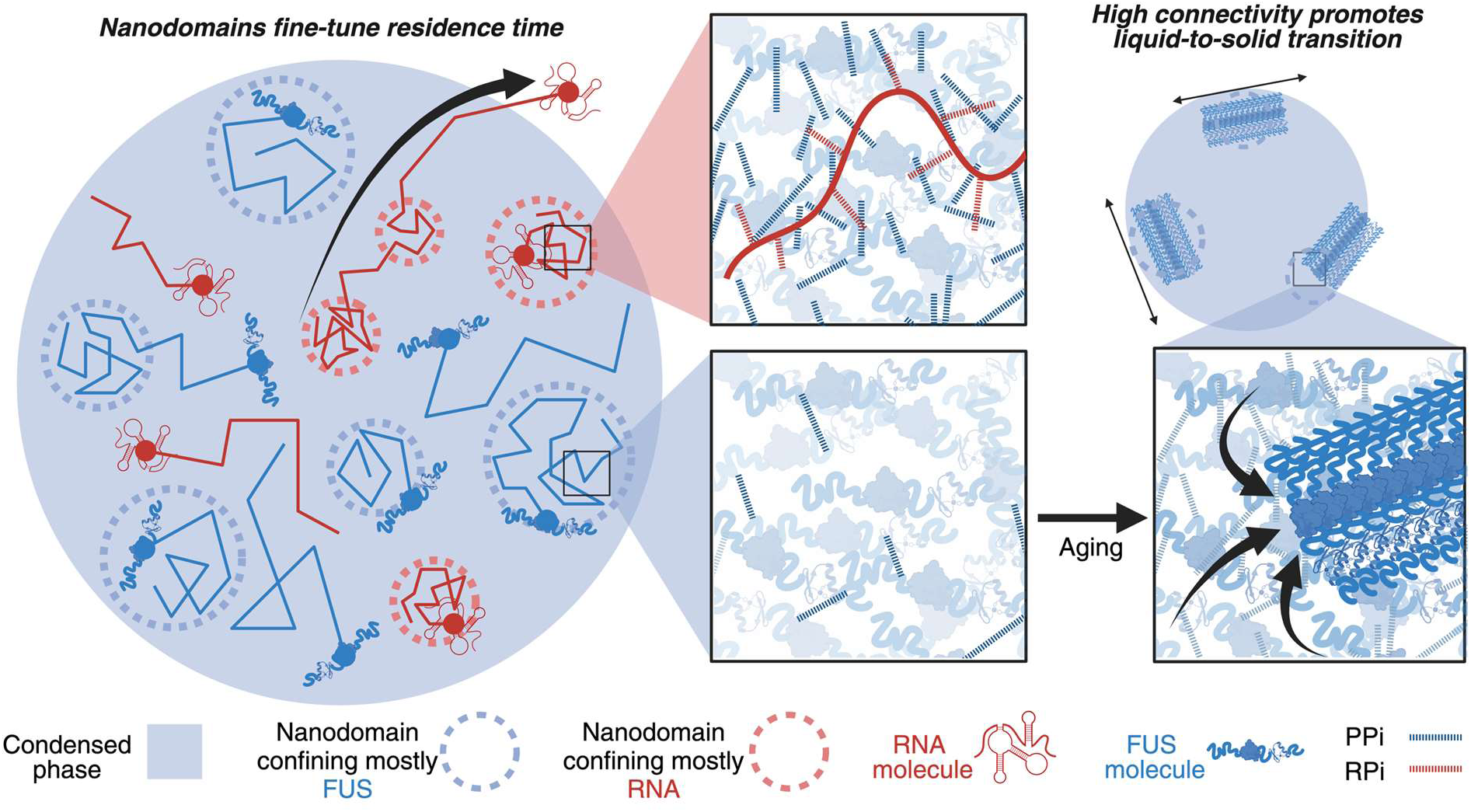
Global condensate functions are determined by nanodomain-regulated local diffusion within condensates. Model showing nanodomains as regions of increased connectivity capable of influencing molecular diffusion and liquid-to-solid transitions within a condensate.

Since nanodomain formation occurs without a change in local condensate density, this regulation of residence time via confined diffusion differs from traditional models that assume normal diffusion within a second, higher density phase. This distinction underscores the critical role of nanodomain dynamics in shaping the fate of individual RNA molecules. The discovery of distinct RNA and protein nanometer-scale domains formed through local percolation, without a density transition, opens new avenues for investigating and clinically leveraging the biophysical mechanisms governing condensate function. Our work therefore has broad implications for modulating biomolecular condensates, for example, using small drug-like molecules, and our ability to monitor the resulting biophysical and pathological behavior.

## Online content

Any methods, additional references, Nature Portfolio reporting summaries, source data, extended data, supplementary information, acknowledgements, peer review information; details of author contributions and competing interests; and statements of data and code availability are available at …

## Methods

### Purification and labeling of FUS and RNA

Full-length tag-free human FUS was purified following a recently developed protocol^25,26^ with a series of ion-exchange chromatography under denaturing condition (Fig. S1), followed by a slow two-step refolding into a storage buffer containing *β*-cyclodextran (BCD) that prevents phase separation. FUS was over-expressed in E. coli BL21(DE3) from pET-FUS plasmid, a generous gift from the Ishihama group. Culture was grown to OD600=1.0 at 37 °C and induced with 1 mM isopropyl β-D-1-thiogalactopyranoside (IPTG). After 6 h of growth at 37 °C, cells were harvested by centrifugation and suspended with buffer A, 10 % glycerol, 20 mM [4-(2-hydroxyethyl)-1-piperazineethanesulfonic acid] (HEPES)-NaOH pH 7.0, 300mM NaCl, 1 mM dithiothreitol (DTT), 1 mM ethylenediaminetetraacetic acid (EDTA), 0.1 % Tween-20, and 1x cOmplete protease inhibitor cocktail (Millipore Sigma), and store for 30 minutes on ice. Cell lysate was sonicated on ice, and insoluble protein was collected by centrifugation. The pellet was solubilized in a buffer B, 6 M Urea, 10 % glycerol, 20 mM HEPES-NaOH pH 7.0, 1 mM DTT, 1 mM EDTA, and 1x cOmplete protease inhibitor cocktail (Millipore Sigma). After centrifugation, supernatant was filtered with 0.45 *μ*m filter (Fisher) and loaded to HiTrap Q column (Cytiva) on a NGC Fast Protein Liquid Chromatography (FPLC) station (Bio-Rad). The flow-through fraction was mixed with CM Sepharose FF resin (GE HealthCare Lifesciences) on a rotator at 4 °C over night and passed through an open column to remove resin. The flow-through fraction was loaded to HiTrap Capto S column (Cytiva) on NGC FPLC station. The flow-through fraction was loaded to HiTrap SP HP column (Cytiva) on NGC FPLC station, and bound proteins were fractionated with 0 to 500 mM linear gradient of NaCl in buffer C, 6 M Urea, 10 % glycerol, 20 mM HEPES-NaOH pH 7.0, 1 mM DTT and 1 mM EDTA. FUS fractions were examined on sodium dodecyl sulfate (SDS) - polyacrylamide gel electrophoresis (PAGE) to ensure purity. If purity was not desired, FUS fractions were pooled, diluted to reduce salt concentration, and re-ran the above protocol from CM Sepharose FF to HiTrap SP HP. Finally, FUS fractions with desired purity (Fig. S1) were pooled, aliquoted to 0.5 mL, flash frozen in liquid nitrogen, and stored in −80 °C freezer until refolding. For the first step in two-step refolding, 0.5 mL denatured FUS was thawed and diluted with 2 mL refolding buffer, 900 mM arginine, 100 mM [N-cyclohexyl-2-hydroxyl-3-aminopropanesulfonic acid] (CAPSO) pH 9.5, 0.3 mM reduced glutathione, 0.03 mM oxidized glutathione, and 1mM ZnCl_2_, and stored over night at room temperature. The second step of refolding was performed during two rounds of two-hours dialysis against buffer D,10 % glycerol, 20 mM HEPES-NaOH pH 6.8, 300 mM NaCl, 0.1 mM EDTA, and 10mM BCD, with a SnakeSkin 10K MWCO dialysis tubing (Fisher) at room temperature. After concentrated with an Amicon 10K MWCO Ultra-4 centrifugal filter unit (Millipore Sigma), the protein concentration of the final product was measured on NanoDrop (Thermo Fisher) with an extinction coefficient of 70,140 M^-1^cm^-1^. The full-length tag-free FUS was aliquoted, flash frozen, and stored at −80°C.

To generate Alexa Fluor 488-labeled full-length tag-free FUS, a third round of dialysis was performed using a buffer D, pH adjusted to 7.5, right after the first two round of dialysis. Alexa Fluor Dye 488 NHS Ester (Click Chemistry Tools) was added to FUS at a dye:protein molar ratio of 20:1 and incubated in dark at room temperature for 30 minutes. Ten rounds of buffer exchange to buffer D were performed using the above filter unit to ensure a complete removal of free dyes. Absorbance spectrum measured on NanoDrop was used to determine the labeling ratio and calculate the effective concentration of Alexa Fluor 488-labeled full-length tag-free FUS.

Alexa Flour 647-labeled firefly luciferase (FL) mRNA was synthesized according to an established protocol^27,52^. A T7 promoter sequence (5’-TAATACGACTCACTATAGGG-3’) was incorporated at the 5’ end of the FL open reading frame during PCR amplification from a pRL-CMV vector (Promega). A Kozak consensus sequence and a 50-nucleotide upstream region was incorporated before the translation start site to ensure enough space for the assembly of translation initiation complex^53^. After *in vitro* transcription with home-made T7 RNA polymerase, the transcript was purified via denaturing PAGE containing 7 M urea. The isolated RNA underwent enzymatic capping (NEB) and polyadenylation (Thermo Fisher), employing an initial incorporation of 2’-azido-2’-dATP (Jena Biosciences) followed by rATP^27,52^. Finally, a click-chemistry reaction with Alexa Flour 647 sDIBO alkyne (Thermo Fisher) was conducted, strategically labeling between the coding sequence and the non-modified poly(A) tail to produce functional, fluorophore-conjugated FL mRNA. Free dye was removed by multiple rounds of washing of the ethanol-precipitated mRNA pellet, which was confirmed by denaturing PAGE (Fig. S1).

### FUS condensate assembly

A 10x phase separation buffer containing 200 mM tris(hydroxymethyl)aminomethane (Tris)-HCl, pH 7.5, 1000 mM NaCl, 20 mM DTT, and 10 mM MgCl_2_ was prepared and filtered with a 0.22 *μ*m syringe filter (Millipore Sigma) for all reconstitution experiments, which was adapted from prior phase separation studies of FUS. Condensates were reconstituted by thoroughly mixing the full-length tag-free FUS at a final concentration of 10 µM, a physiological concentration of FUS in Hela cell nucleus^13^, with the phase separation buffer, an optional OSS, and any fluorescently labeled species based on experimental type #1-3 (Fig. 1h), as described below. The time from assembly to data acquisition was kept under 30 minutes unless specified otherwise.

For condensate SMT experiments, samples were prepared with 10 nM Alexa Fluor 488-labeled FUS and 50 pM of various fluorescent species in phase separation buffer. An oxygen scavenging system (OSS) consisting of PCA-PCD-Trolox was used for boundary detection experiments, while a GODCAT OSS was employed for extended trajectory reconstruction measurements to reduce autofluorescence. For FUS SMT without boundary detection, 50 pM Alexa Fluor 488-labeled FUS was used. Dual-color trajectory reconstructions included both labeled FUS and RNA. OSS was omitted for fast-diffusing particles with low signal-to-noise ratios. When applicable, 10% (w/v) Dextran T-500 and 50 ng/μL Human HeLa Cell total RNA were added as crowding agent and total RNA, respectively.

FUS condensates were prepared with modifications for various experiments. For phase diagram studies (Fig. S2), a 10x phase separation buffer without NaCl and MgCl2 was used to control salt and FUS concentrations. Rheology measurements (Fig. 1c) employed a C-Trap optical tweezer with confocal microscope, using 200 nM Alexa Fluor 488-labeled FUS and PCA-PCD-Trolox OSS for adequate SNR. Z-stack imaging (Fig. 1g) utilized an Alba5 confocal microscope with 40 nM Cy5 free dye to visualize 3D shape and glass slide location. 40 µM of Edaravone^54^ and 10µM of Riluzole^46,55^ were added to condensates after drugs where solubilized in DMSO. A 40 µM DMSO control with no dissolved drug was used as a control (Fig. 5). Fibrilization imaging (Fig. 4a, 5b, S18, S19) involved FUS condensates assembled without OSS, crowding reagents, or fluorescent species to exclude potential artifacts.

### Microrheology

Condensate fusion events were measured in a u-Flux microfluidics chamber (LUMICKS). The chamber was thoroughly cleaned with water, bleach, sodium thiosulfate, BSA, and Pluronic acid F-127 (Millipore Sigma) to prevent unwanted interactions. Experiments used two channels: a condensate channel and a measurement channel. Condensates were pumped into the first channel at low pressure (<0.5 bar), while the measurement channel contained 400 nM FUS (saturated concentration from phase diagram, Fig. S2). For microrheology measurements (Fig. 1c), ∼1 μm diameter condensates were selected. The probing trap approached the measurement trap at 50 nm/s, with force measured at 50 kHz and confocal imaging at 4 Hz. This setup allowed for precise control and measurement of condensate fusion events.

Analysis of force traces and confocal imaging were performed with home-made python scripts and Pylake package (LUMICKS). All home-made python scripts used in this section and below were all deposited at https://github.com/walterlab-um/intra_condensate_SPT. For confocal imaging data, condensate area was calculated by Otsu thresholding on fluorescence intensity, which has been proved to be the most accurate condensate boundary estimator among other computer vision methods for large condensates^56^. For force data, fusion events were identified by down sampling from 50 kHz to 5 Hz, calculating the gradient of force, and finding the largest peak in the gradient of force. For each fusion event, the original 50 kHz force data (*F*) was fitted to a previously used model^13,28^:

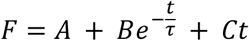

where *τ* is the characteristic fusion time and *A*, *B*, *C* are constants. The exponential part is a theoretical formula for droplet fusion. Constant A describes background of force measurement while constant C describe the artificial force increase introduced by the interference between optical traps when probing trap approaches measurement trap at a constant speed. All fusion events were manually inspected before subjected to the above pipeline to exclude traces generated by non-fusing condensates or debris.

#### SMT via HILO microscopy

FUS condensates were tethered to glass slides using a PEG-biotin:streptavidin:FUS-biotin system we developed. Glass coverslips were functionalized with APTES, mPEG-SVA, and biotin-PEG-SVA (1000:1 ratio), followed by DST treatment^57^. Sample wells were created using half-cut PCR tubes (Sigma-Aldrich) glued to the prepared slides with epoxy (Ellsworth Adhesives) (Fig. S20), and the tethering system was established with 1 mg/mL streptavidin and 1 μM biotinylated FUS to prevent translational and rotational motions (Fig. S21). Mineral oil was added to prevent evaporation. Imaging was performed on an Oxford Nanoimager S with a 100x 1.4 NA oil immersion super apochromatic objective, 405, 473, 532, 640 nm lasers with appropriate dual-band emission filters, and a Hamamatsu sCMOS Orca flash 4 V3 camera, using HILO microscopy^58^ (52° laser angle) at 24 °C with 117 nm pixel size^59^. Three main imaging protocols were employed (Fig. 1h): (1) intra-condensate SMT/SPT with boundary detection, using red (640 nm, ∼30 mW) and green (473 nm, ∼2 mW) channels; (2) intra-condensate FUS SMT without boundary detection, using only the green channel (473 nm, ∼30 mW); and (3) dual-color trajectory reconstructions, using alternating-laser excitation of 473 nm and 647 nm lasers. For FL mRNA SMT outside condensates, FUS concentration was reduced to 1 μM, and the focal plane was positioned above condensates.

### Phase diagram and fibrilization imaging

Phase diagram samples were prepared at various FUS concentrations of 0.5, 1.0, 2.0, and 5.0 µM. At each concentration we systematically varied the sodium chloride or potassium chloride salt concentrations of 0.1, 0.5, 0.75, and 1.0 M. For an accurate measurement for condensation near phase boundary of the phase diagram, an incubation time of one hour was used to give the protein -condensates adequate time to coarsen into a detectable size. Samples were imaged on an Olympus IX81 microscope equipped with a 100x oil-immersion objective with ∼1 mW of 488 nm laser and an exposure time of 100 ms. Fibrilization imaging was performed in bright field at various time points of 0, 8, and 24 hours using the Nanoimager (Oxford) described above with a 100 ms exposure time.

### Diffusion profiling pipeline

To pick up only single particle fluorescence signal while suppressing background, band-pass filtering on all SMT/SPT videos was achieved by applying a difference of Gaussian (DoG) filter with a lower sigma of 1 pixel and a higher sigma of 3 pixels, representing on-focus and slightly off-focus single molecules. Filtered videos were subjected to TrackMate^60^ to detect spots and extract trajectories. Spot detection was performed with a Laplacian of Gaussian (LoG) detector, where an estimated object diameter was set to 5 pixels (585 nm) and a spot quality threshold set to 5 or determined by the peak of false-positive spots in the distribution of quality values (Fig. S22). Trajectories were extracted using the Linear Assignment Problem (LAP) algorithm with specific settings for intra-condensate tracking (maximum linking distance: 5 pixels, no gaps) and dilute-phase tracking (maximum linking distance: 15 pixels, no gaps). Exported trajectories were analyzed using custom Python scripts for diffusion profiling.

The diffusion profiling pipeline consists of discrete categorization and continuous spectrum analysis (Fig. 2a-f). Trajectories were classified as immobile (<30 nm mean step size), confined (*⍺* < 0.7)^33^, or normal diffusion (*⍺* > 0.7) based on mean step size and anomalous diffusion component *⍺*. The threshold of 30 nm was based on the 16 nm static localization error calculated from static mRNA molecules attached to glass surface (Fig. S23). The *⍺* value was calculated by fitting the (MSD)-lag time (*τ*) curve on a log-log scale with the formula below.

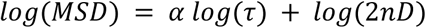

Here, *n* is the dimension of tracking (*n* = 2 in our case) and *D* is the diffusion coefficient. Half the trajectory length was used for optimal fitting^61^ unless it’s shorter than 5 steps, where a minimal number of 4 MSD-*τ* points were guaranteed for accurate fitting. Trajectories with R² < 0.7 including trajectories with extremely small α (Fig. S24) were excluded.

The continuous spectrum analysis generated distributions of mean step size, *α*, angle between steps, apparent diffusion coefficient (*D*_*app*_), and localization error. Mean step size was calculated for all trajectories, while α components were determined only for mobile trajectories. The angle between steps, ranging from 0° (no direction change) to 180° (complete reversal), was calculated for all mobile trajectories, regardless of turn direction. The *D*_*app*_ and localization error were calculated for non-confined trajectories by fitting MSD-*τ* curve to the following formula, which is optimized for least-square fit of MSD^61,62^:

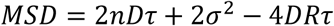

where *σ* is the localization error and *R* is the motion blur coefficient (*R* = 1⁄6 for continuous imaging as in our case). A Bayesian statistics-based state array method (saSPT^34^) was used to generate higher-resolution *D*_*app*_ spectrum analysis.

### Trajectory reconstruction and step size heatmap analysis

Point accumulation reconstruction was performed with home-made python scripts on 10,000-frames long SMT videos collected from trajectory reconstruction experiments (Fig. 1h, #2-3) as described above. We ensured sufficient coverage of locations within a condensate to avoid pseudo-puncta from a few trajectories by introducing an oxygen scavenging system (OSS) that extended SMT video length from the previous 200 frames, 4 s to 10,000 frames, 200 s (Fig. 1h, Fig. S4). To ensure that puncta in trajectory reconstruction images reveal underlying intra-condensate architecture that confines diffusion, the oversampling from long trajectories needs to be suppressed, avoiding the formation of artificial puncta due to a single long trajectory. Therefore, when generating the pool of locations for the reconstruction, only 10 locations were randomly sampled from all locations for any trajectory longer than 10 steps while all locations from trajectories shorter than 10 steps were passed along. trajectory reconstruction image of the whole FOV was rendered by binning the pool of locations back to a grid of 117-nm pixels, the same as the input trajectory reconstruction video. Similarly, step size heatmap image of the whole FOV was generated from the pool above, by calculating pixel-by-pixel the mean for all steps that have a center of step within the pixel. Drift correction was not needed given the short total length of imaging time (200 s).

Not all condensates were subjected to downstream analysis because some condensates did not have enough trajectories to cover the whole condensate while others had a strip-like pattern due to ‘lensing effect’, where the spherical shape of condensates works as a lens to skew illumination (Fig. S25). Therefore, qualified condensates were picked by a python script followed by manual inspection. The python script first generates contours for all condensates by applying a threshold of 10 locations per pixel to the whole-FOV trajectory reconstruction image smoothed with a *σ* = 1 Gaussian kernel. The whole-FOV trajectory reconstruction image was split to individual-condensate trajectory reconstruction images by filtering out condensates smaller than 200 pixels^2^ and cropping out each remaining condensate with a surrounding box with a padding distance of 3 pixels. All individual-condensate trajectory reconstruction images were manually inspected to leave out all condensates with unwanted strip-like pattern (Fig. S25). Trajectory reconstruction and step size heatmap images of all qualified condensates were paired, such as trajectory reconstruction images of FUS versus mRNA, step size heatmap of FUS versus mRNA, and trajectory reconstruction versus step size heatmap images of FUS, and analyzed with weighted pixel-wise Pearson correlation coefficient (PCC) using the below formula:

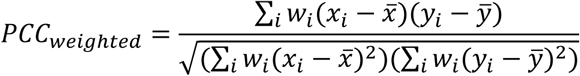

where *x*_*i*_ and *y*_*i*_ are the value of *i*-th pixel in the vectorized (*X*, *Y*) image pair, *x̅* and *ȳ* are the mean of all pixels in the *X* or *Y* image, and *w*_*i*_ is the weight of *i*-th pixel in the same vectorized image. *X* and *Y* image in the (*X*, *Y*) image pair can be either trajectory reconstruction image, where pixel value is number of locations, or step size heatmap image, where pixel value is mean step size. Weight is set to be the square of number of locations per pixel, factoring in that pixel values coming from more trajectories are more reliable than those coming from less trajectories.

### Pair-correlation functions

spatial distribution of SMT trajectories within a condensate was analyzed by the auto- or cross-pair correlation function *G*_*auto*_(*r*) or *G*_*cross*_(*r*) (a.k.a. radial distribution function) of locations. Pair correlation function *G*(*r*) describes the probability of finding a particle of interest at a distance *r* from a reference particle, where the reference particle and the particle of interest are of same type in *G*_*auto*_(*r*) but are of different type in *G*_*cross*_(*r*). pair correlation function can be calculated by

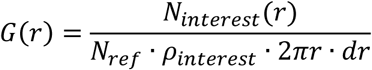

where function *N*_*interest*_(*r*) counts the number of particles of interest that are *r* to *r* + *dr* away from reference particles in a condensate, *N*_*ref*_ is the total number of reference particles in a condensate, *ρ*_*interest*_ is the density of particles of interest in a condensate, and 2*πr* · *dr* is the ring area that normalizes the excess number of particles counted by function *N*_*interest*_(*r*) as the ring area scales with *r*.

In the python script, *N*_*interest*_(*r*) was calculated by iterating through all reference particles because the normalization factor needs to be adjusted when the reference particle was too close to the condensate boundary. In that case, the ring centered at the reference particle was partially outside the condensate, where no particles of interest can be found, and thus it would be wrong to assume an average number of counts of *ρ*_*interest*_ · 2*πr* · *dr* can be found in the ring. Therefore, for all reference particles closer to the condensate boundary than *r* + *dr*, the ring area within condensate was calculated to substitute 2*πr* · *dr* in the above equation. For all *G*(*r*) in this study, *dr* was kept at 100 nm and *r* ranged from 0 to 1 *μ*m, with a sliding-window of 20 nm between adjacent *r*.

Every qualified condensate yielded a *G*(*r*) and the *G*(*r*) for all plots was a weighted average of all the individual-condensate *G*(*r*), with the weight being total number of particles within each condensate. Each individual-condensate *G*(*r*) was then fitted with the exponential model below because exponential decay can depict the behavior of molecules in a phase-transition system^84^:

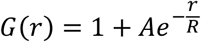

where *A* is the amplitude that describes how much more or less likely a particle could be found compared to a random distribution of particles, and *R* is the characteristic radius of the clustering of particles. A parametric bootstrapping, 5,000 rounds of re-sampling from the pool of individual-condensate *G*(*r*), was performed to estimate the *A* and *R* for each experiment and their confidence intervals. *A* and *R* in each bootstrapping round were also plotted as histograms.

### Spatial distributions of nanodomains

Nanodomains locations were estimated by either puncta detected by a LoG spot detector on individual-condensate trajectory reconstruction images or using the centroid of confined FUS or RNA trajectories within a condensate. The distance of all such locations within a condensate to the center of the condensate was normalized by the radius of the condensate, where the radius was calculated from the total area of the condensate assuming a circular shape. Like *G*(*r*), the distribution of all such distances within a condensate was also biased by the ring area growing with *r*, and thus needed to be normalized by ring area using:

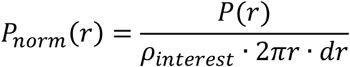

where *P*(*r*) and *P*_*norm*_(*r*) are the probability and the normalized probability of distances that are *r* to *r* + *dr* away from the condensate center, *ρ*_*interest*_ is the density of particles of interest in the condensate, and 2*πr* · *dr* is the ring area. Every qualified condensate yielded a *P*_*norm*_(*r*), and 5,000 rounds of non-parametric bootstrapping from the pool of individual-condensate *P*_*norm*_(*r*) were performed to estimate the mean of *P*_*norm*_(*r*) for every *r*.

### Fluorescence lifetime imaging and correlation spectroscopy

Fluorescence lifetime imaging (FLIM) imaging of tethered condensates was performed with an Alba5 time-resolved laser-scanning confocal microscope (ISS, Inc) with a pulsed supercontinuum broadband laser excitation source (Fianium WhiteLase SC-400-8-PP), avalanche photodiode (APD) detectors, and a Beckr-Hickl SPC-830 TCSPC module with a 531/40nm filter. 488 nm excitation was selected using acousto-optic tunable filters and ∼5uW average power was selected so that the mean counts per second was less than approximately 100kcps to avoid dead-time artifacts in the fluorescence lifetime fits. Condensates tethered on the glass surface were first found under regular scanning confocal imaging mode. After an optimal field of view framing a single condensate was identified, ten rounds of FLIM scanning throughout the whole field of view were performed to ensure adequate data points for the lifetime calculation via deconvolution and fitting to a reference instrument response function (IRF) using FLIMfit^63^. The number of FLIM scans on each condensate was carefully chosen to achieve a total imaging time of around five minutes, matching a time scale where nanodomains were relatively immobile as revealed by time-lapse trajectory reconstruction.

The coupled fluorescence correlation spectroscopy (FCS) with FLIM assay was performed with the same setup as above, but only ten locations within each condensate were chosen at random using Sobel sampling rather than scanning through the whole condensate as in FLIM imaging. One-minute-long time-resolved intensity traces for Alexa488-labeled full-length tag-free FUS were acquired per location and analyzed both for diffusion coefficient using FCS as well as fluorescent lifetime. Lifetime fitting was done with a MATLAB package FluoFit^64^, while FCS curves were analyzed with a custom MATLAB implementation of a segmented autocorrelation algorithm^65^ to reduce the effect of photobleaching or long-term diffusive effects such as rocking of the condensate droplet as a whole. Note that the one-minute-long FCS traces provide much more datapoints for FLIM fitting compared to each pixel in a FLIM image, resulting in a better fit and explaining the different ranges of lifetime in FLIM imaging versus FCS-FLIM assays.

## Acknowledgement

We sincerely thank Sujay Ray, Ziyuan Chen, Adam Decker, Natalie Rogers, and Xiaofeng Dai for their insightful discussions on developing the analysis pipelines for our SMT datasets. We appreciate help from Damon Hoff at the Single Molecule Analysis in Real-Time (SMART) Center of Biophysics at the University of Michigan, for FLIM and FCS measurements and analysis, and Ehsan Akbari and Denis Pelekhov at the Nano-Systems Laboratory (NSL) of the Physics department at the Ohio State University, for their help in using a LUMICKS C-Trap for condensate microrheology measurements. We much appreciate the invaluable feedback and proofreading efforts by Ziyuan Chen, Adrien Chauvier, Minjun Jin, Alexander Johnson-Buck, Sicong Ma, Yi Zhu, and the University of Michigan U-M GPT as well as ChatGPT. We also thank Liuhan Dai for his early joint efforts on protein purifications and Mohammed Hijaz for efforts on quantifying the translatability of FL mRNAs. N.G.W. acknowledges funding from NIH grant R35 GM131922, a sub-award of NIH grant R01 NS097542, and Chen-Zuckerberg Initiative (CZI) grant 2022-250725; whereas E.R.S. is thankful for an NSF GRFP fellowship DGE2241144.

## Author contributions

All authors contributed substantially to discussion of the content and writing of the article. G.G., E.R.S. and N.G.W. reviewed and edited the manuscript before submission. G.G. and E.R.S. performed the bulk of the experiments and analysis for the article.

## Competing interests

The authors declare no competing interests.

## Additional information

Extended data for this paper are available at…

## Notes

### Competing Interest Statement

The authors have declared no competing interest.

### Summary of Updates

Numerous experiments were added and the manuscript thoroughly rewritten.

