## Supplementary Figures and Methods for "Nanoscale domains govern local diffusion and aging within FUS condensates"

| *Target* | Immobile Fraction | Confined Fraction | Normal Diffusion Fraction | Mean Step Size (nm) | Mean anomalous component α | Mean angle between steps (degree) | Major Peak of $D_{app}$ ($\mu m^2/s$) |
|---|---|---|---|---|---|---|---|
| *FUS-Alexa Fluor 488, condensed phase* | 0.17 ± 0.06% | 67 ± 1% | 33 ± 1% | 172.9 ± 0.2 | 0.818 ± 0.001 | 105.44 ± 0.05 | 0.811 |
| *FL mRNA-Alexa Fluor 647, condensed phase* | 6 ± 1% | 83 ± 2% | 11 ± 2% | 60.9 ± 0.2 | 0.613 ± 0.002 | 116.44 ± 0.05 | 0.076 |
| *FL mRNA-Alexa Fluor 647, dilute phase* | 6 ± 3% | 40 ± 2% | 54 ± 5% | 810 ± 3 | 1.023 ± 0.005 | 93.5 ± 0.3 | 10.975 |
| *miRNA-21 -Cy5, condensed phase* | 0.32 ± 0.08% | 46 ± 2% | 54 ± 2% | 153.02 ± 0.09 | 0.8881 ± 0.0006 | 97.86 ± 0.03 | 0.433 |
| *Cy5 dye, condensed phase* | 1.8 ± 0.5% | 53 ± 1% | 45 ± 1% | 228 ± 1 | 0.889 ± 0.003 | 98.9 ± 0.1 | 2.848 |
| *20-nm beads, condensed phase* | 45 ± 9% | 40 ± 6% | 15 ± 4% | 47.0 ± 0.4 | 0.708 ± 0.005 | 104.1 ± 0.1 | 0.031 |
| *20-nm beads, dilute phase* | 0 ± 0% | 35.6 ± 0.6% | 64.4 ± 0.6% | 750 ± 3 | 0.980 ± 0.006 | 91.9 ± 0.3 | 9.006 |
| *200-nm beads, dilute phase* | 0.5 ± 0.5% | 23.3 ± 0.6% | 76 ± 1% | 340 ± 2 | 1.039 ±0.005 | 82.9 ± 0.3 | 1.874 |
| *$QD_{neg}$, condensed phase* | 23 ± 4% | 57 ± 5% | 21 ± 1% | 51.12 ± 0.09 | 0.743 ± 0.001 | 100.23 ± 0.04 | 0.028 |
| *$QD_{neu}$, condensed phase* | 66 ± 5% | 25 ± 5% | 9.6 ± 0.3% | 48.93 ± 0.07 | 0.747 ± 0.001 | 104.74 ± 0.03 | 0.035 |
| *$QD_{pos}$, condensed phase* | 20 ± 2% | 55 ± 3% | 25 ± 3% | 52.03 ± 0.06 | 0.7441 ± 0.0008 | 100.47 ± 0.02 | 0.028 |

**Table. S1 │ Metrics calculated from the diffusion-profiling pipeline.**

The pipeline aims to depict the complete diffusion profile of target molecules or particles, leading to a discrete categorization that divides molecules/particles into discrete diffusion classes via direct thresholding and a continuous spectrum part that focuses on population-wide shifts in mean step size, $\alpha$ component, angle between steps, and $D_{app}$ of normal-diffusion molecules. The table shows the mean ± standard error of mean (SEM) of discrete fractions and each spectrum. Condensed phase rows are shaded.

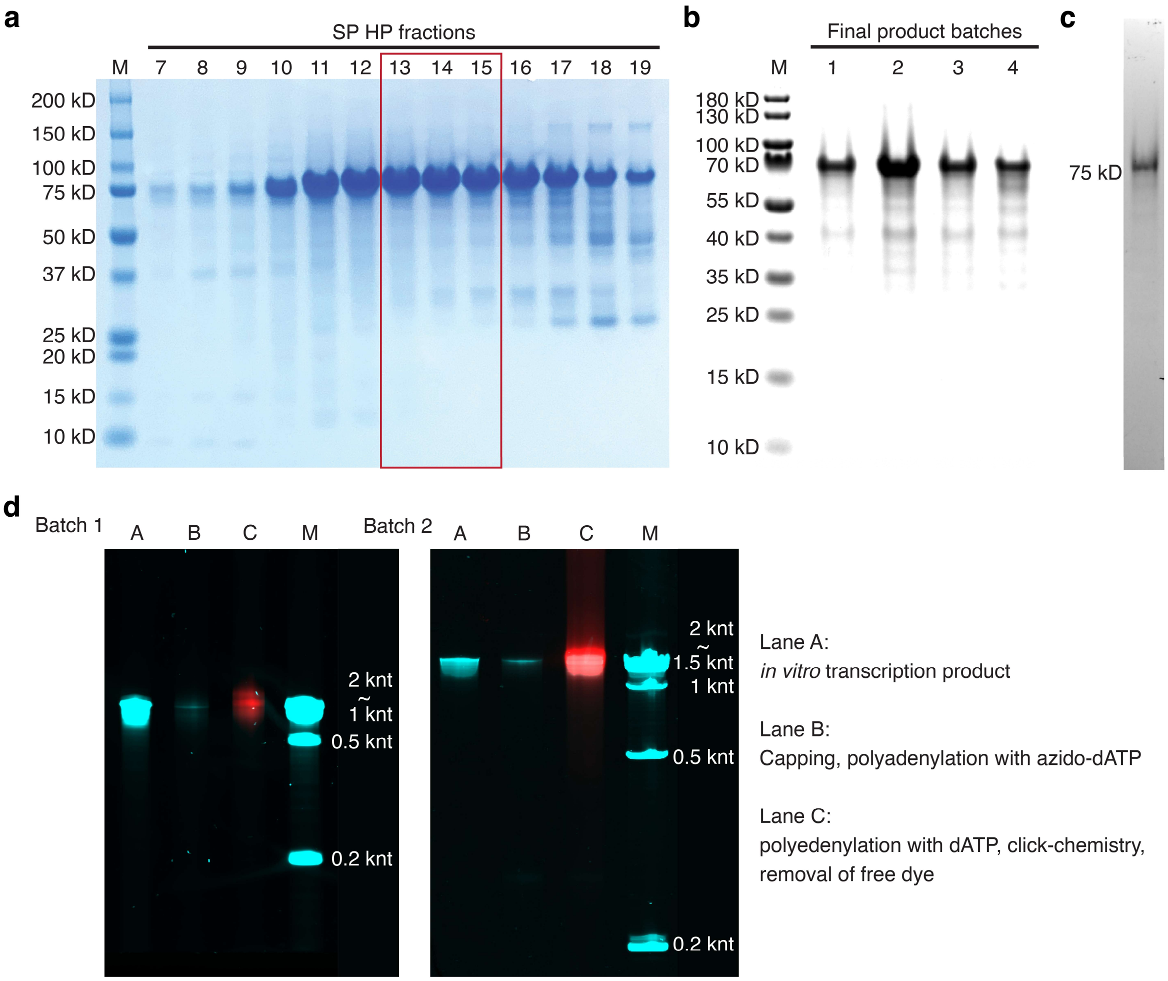


**Fig. S1 | Purification and labeling of human full-length tag-free FUS and firefly luciferase (FL) messenger RNA (mRNA).**

**a |** A sodium dodecyl sulfate (SDS)-polyacrylamide gel electrophoresis (PAGE) gel stained with Coomassie Blue showing fractions from the HiTrap SP HP (Cytiva) cation exchange chromatography, the last chromatography step of the purification of human full-length tag-free Fused-in-Sarcoma (FUS) protein. The label "M" marks the lane of protein molecular weight ruler, PageRuler (Thermo Fisher Scientific). The red frame denotes the only three fractions pooled for subsequent steps to prioritize purity over yield. **b |** Four batches of refolded FUS, the final product of FUS purification, on an SDS-PAGE gel stained with Coomassie Blue. **c |** Alexa Fluor 488-labeled FUS on an SDS-PAGE gel imaged by a Typhoon imager (Cytiva). **d |** Two batches of purification and labeling of FL mRNA analyzed by a 5% 8 M urea PAGE gel stained with SYBR Gold (Thermo Fisher Scientific) and imaged using a Typhoon 5 imager, demonstrating the whole procedure from the PAGE-gel purified *in vitro* transcription product to the capped, polyadenylated (via a between-body-and-tail (BTT) strategy), and Alexa Fluor 647-labeled final product. The SYBR Gold RNA stain signal is pseudo-colored as cyan and the Alexa Fluor 647 signal as red. No shifts are observed from Lane A to Lane C, indicating that the poly(A) tail added to the mRNA is reasonably short and beyond the detection limit of a 5% urea PAGE gel, mimicking the average poly(A) length of ~200 nt in mammalian cells[1]. Note that SYBR Gold staining has a worse detection limit than the Alexa Fluor 647 covalently linked to mRNA so that the SYBR Gold signal may not always be observable in Lane C due to inevitable losses of product during each step.

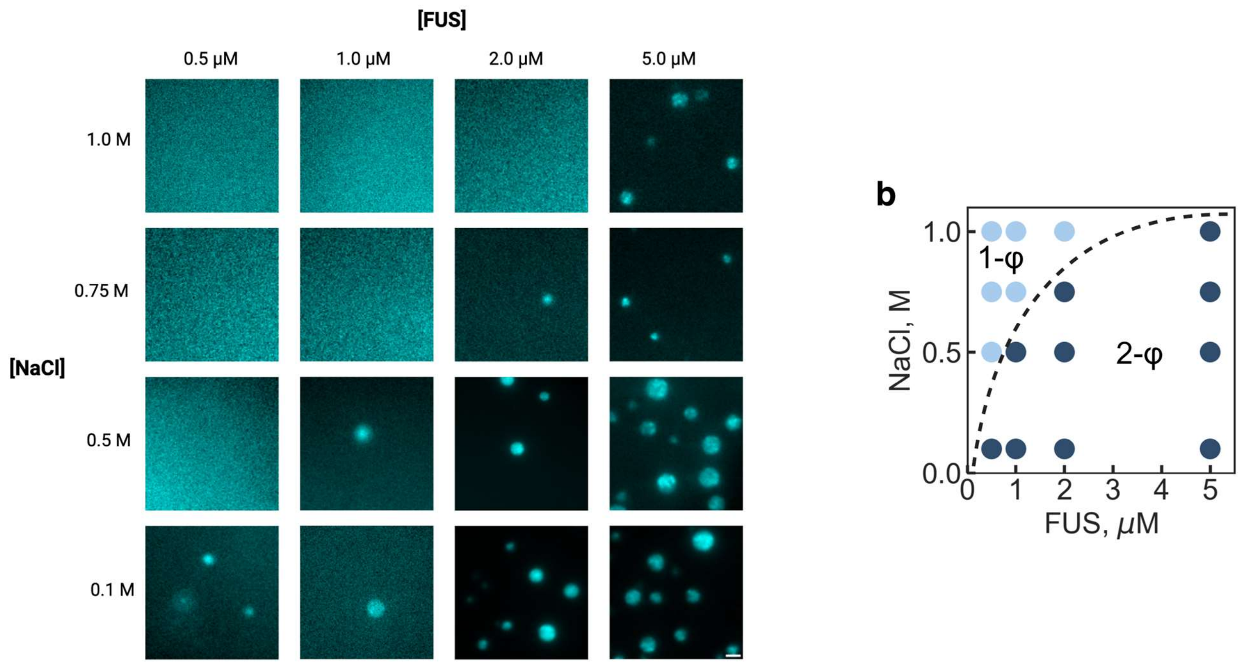


**Fig. S2 │ Phase diagram of full-length tag-free FUS.**

**a │** Representative epi-fluorescence images of FUS condensate assembled with 10 nM Alexa Fluor 488-labeled FUS under the protein and salt concentrations specified on the x- and y-axis. **b │** Phase diagram reconstructed from **a**. The dotted line is a guide for the eye, denoting the phase boundary (or binodal curve). Dark blue dots denote conditions where condensates can be observed (two-phase region, i.e., 2- ϕ) while light blue dots denote conditions where condensates cannot be observed (single-phase region, i.e., 1- ϕ).

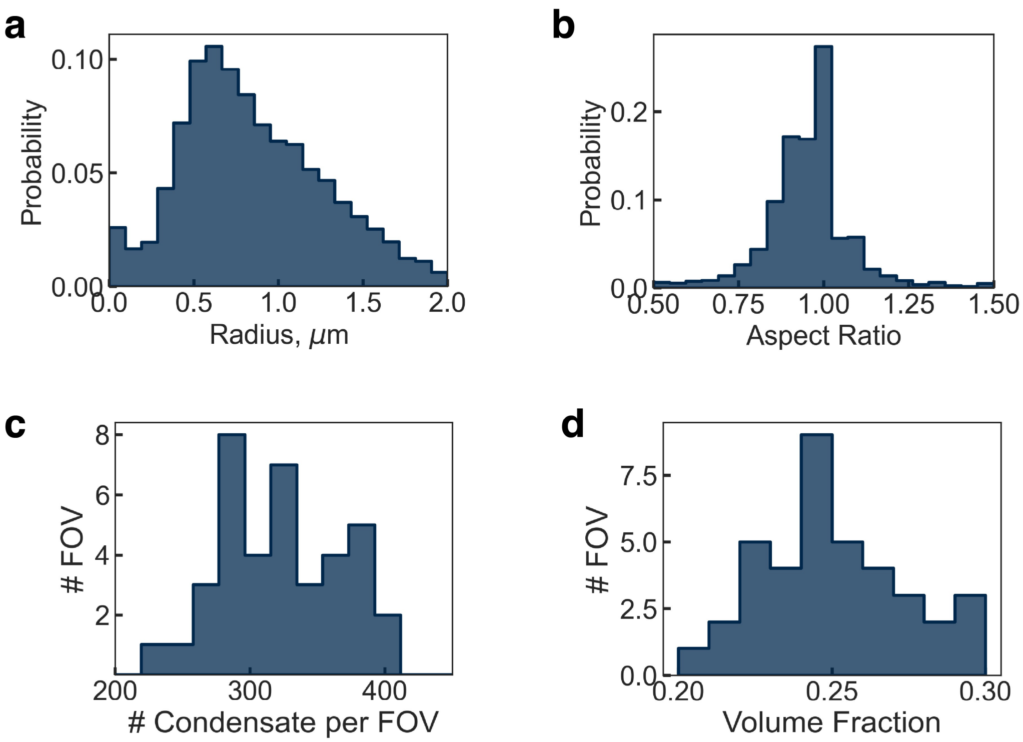


**Fig. S3 │ The properties of tethered FUS condensates.**

**a │** Distribution of condensate size. The equivalent radius of each tethered condensate was calculated from the contour area of the detected condensate boundary, assuming a circular shape. Condensate boundary detection was achieved using a machine-learning-based pixel classifier, which has been identified to be the best condensate boundary detection method for condensates with a broad range of sizes[2]. **b │** Distribution of the aspect ratio of tethered condensates, defined as the ratio between the width and height of a bounding rectangle of the detected condensate boundary. An aspect ratio closer to 1 suggests a more circular condensate, which is a signature morphology of highly liquid-like droplets. **c |** Distribution of the number of condensates per the field of view (FOV) of the microscope, representing the number of condensates simultaneously available for intra-condensate single molecule tracking (SMT) analysis. **d |** Distribution of the volume fraction of condensates per FOV, defined as the total condensate boundary contour area in each FOV divided by the total area of the FOV. This value ensures that our intra-condensate SMT experiments are not performed on condensates over-crowded on a slide.

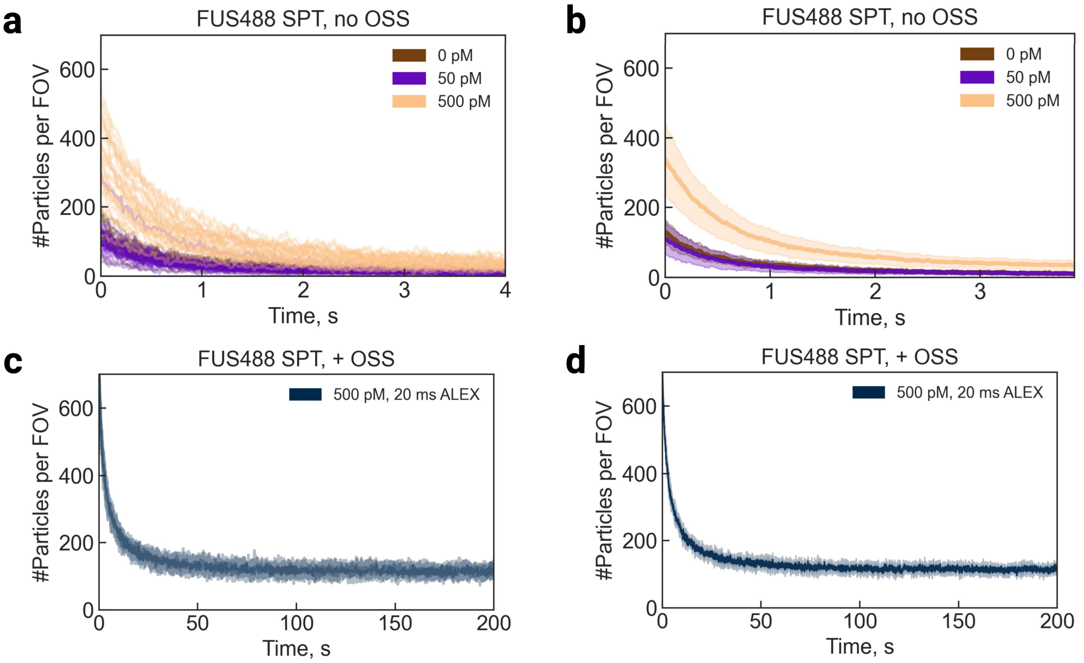


**Fig. S4 | Effect of oxygen scavenging system (OSS) on elevating the number of trackable FUS molecules to ensure an adequate number of trajectories for SMT-PAINT.**

**a** | The change in number of trackable Alexa Fluor 488-labeled FUS molecules in the microscope FOV over 4 s, 200 frames. Each curve is from one SMT video. A comparison of 0, 50, and 500 pM concentrations of labeled FUS shows a concentration-dependence of number of particles, strongly suggesting the SMT trajectories indeed derive from labeled FUS. The decay is caused by photobleaching. The plateau is formed by the replenishment of fluorescent molecules rejoining the highly inclined and laminated optical sheet (HILO)[3] illumination plane as well as the focal plane via axial diffusion. **b** | The mean, shown as a solid line, and the standard deviation (STD), shown as shaded area, of **a**. **c** | The change in number of trackable Alexa Fluor 488-labeled FUS molecules in the microscope FOV over 200 s, 10,000 frames with the addition of coupled glucose oxidase and catalase (GODCAT) OSS[4]. Each curve derives from one SMT video. Note that both the decay has been extended and the plateau has been elevated by the addition of OSS. **d** | The mean, shown as a solid line, and the STD, shown as shaded area, of **c**.

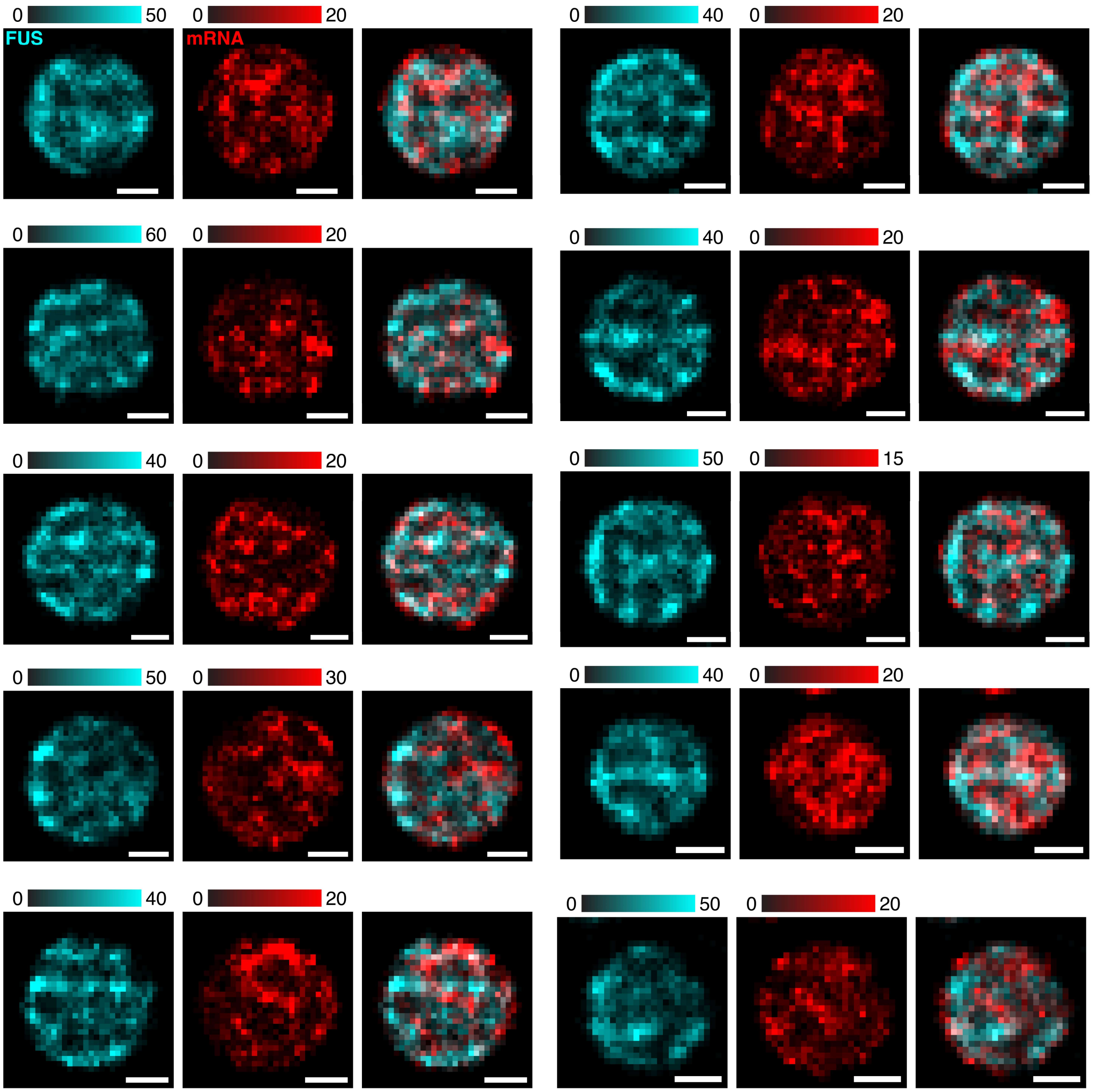


**Fig. S5 │ Representative dual-color SMT-PAINT images of FUS and FL mRNA.**

Color bars are in the unit of number of single-molecule localizations per pixel. All scale bars are 1 μm.

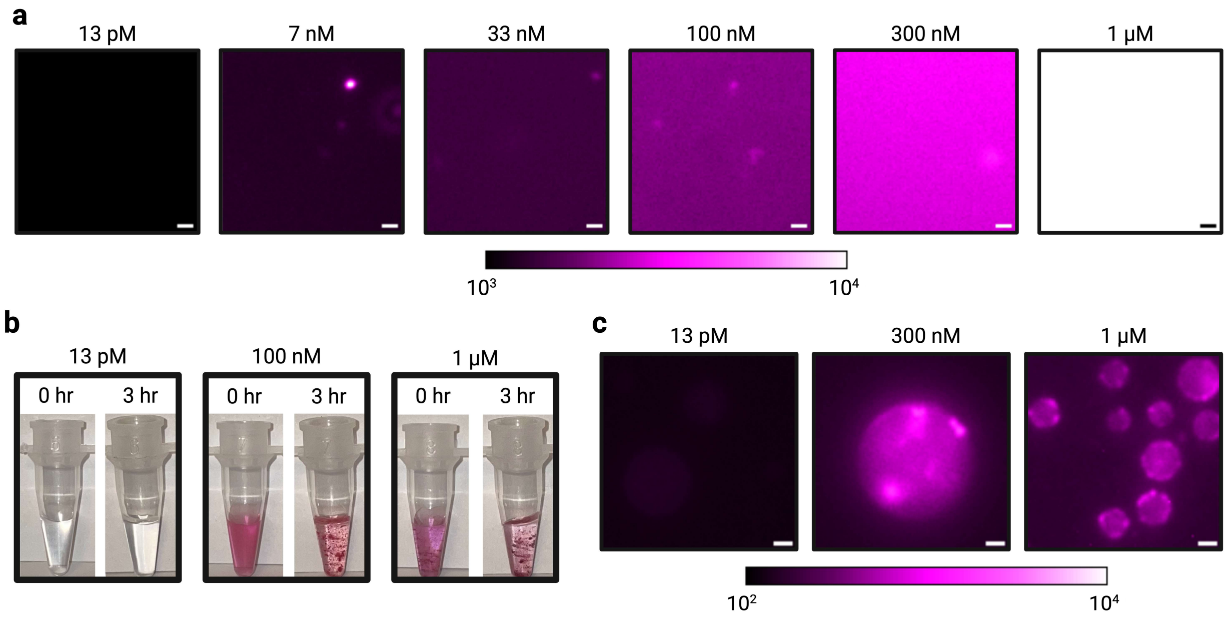


**Fig. S6 | Nile Red puncta within condensates are a result of dye aggregation.**

**a |** Aggregation of Nile Red due to limited solubility in the water-based 1x phase separation buffer at a concentration as low as 7 nM, revealed by HILO fluorescence imaging of a concentration series of Nile Red freshly diluted from a DMSO-based stock solution into water-based 1x phase separation buffer. **b |** Photos of aggregated Nile Red in the water-based 1x phase separation buffer at varying concentration and time. Nile Red aggregates are clearly visible by the naked eye at a concentration as low as 100 nM after incubation for as short as 3 hours. **c |** Nile Red imaging of freshly reconstituted FUS condensates in a series of concentration under HILO illumination. Bright Nile Red puncta are observed within condensates, preferentially located at the condensate peripheries, even without the need of aging, in contrast to previous reports[5,6]. Overall, these results suggest potential limits for using Nile Red as an indicator for the intra-condensate chemical environment.

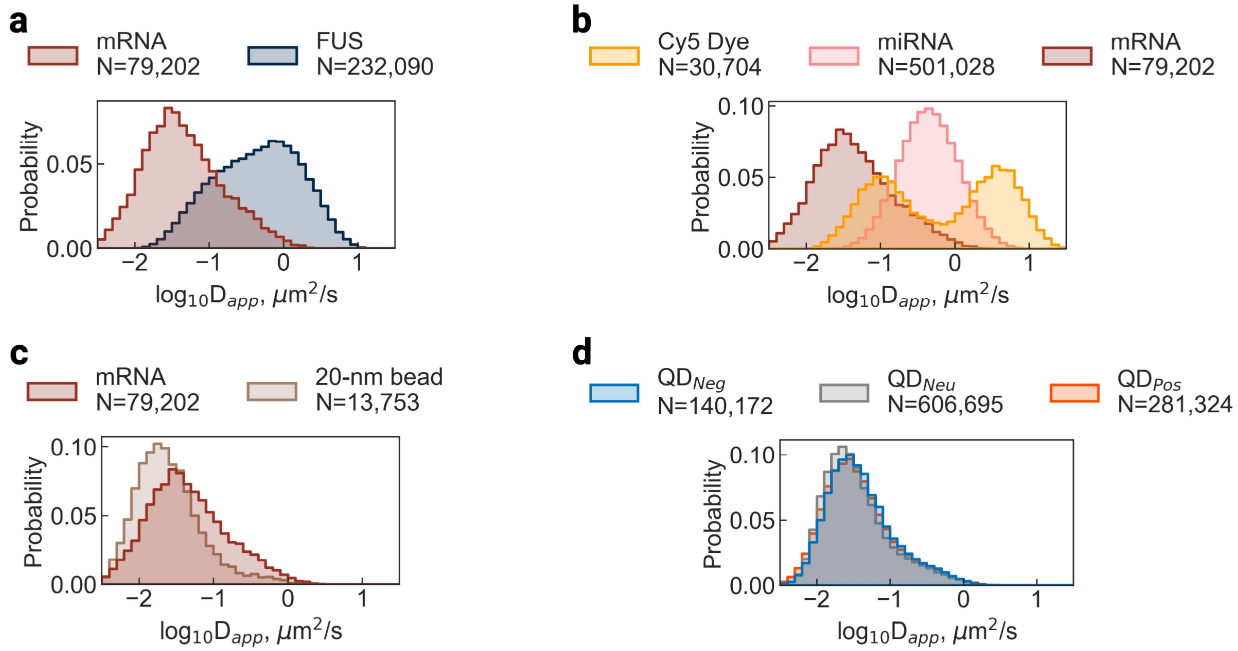


**Fig. S7 | Distributions of apparent diffusion coefficient $D_{app}$ determined by canonical per-trajectory MSD-τ fitting.**

**a-d** | Distributions of $D_{app}$ calculated from an optimized form of MSD-τ fitting[7,8] corresponding to the datasets in Fig. 2b-c (**a**), Fig. 2f-g (**b**), Fig. 2h-i (**c**), and Fig. 2j-k (**d**). Details of the fitting can be found in the *Diffusion profiling pipeline* section under *Methods*.

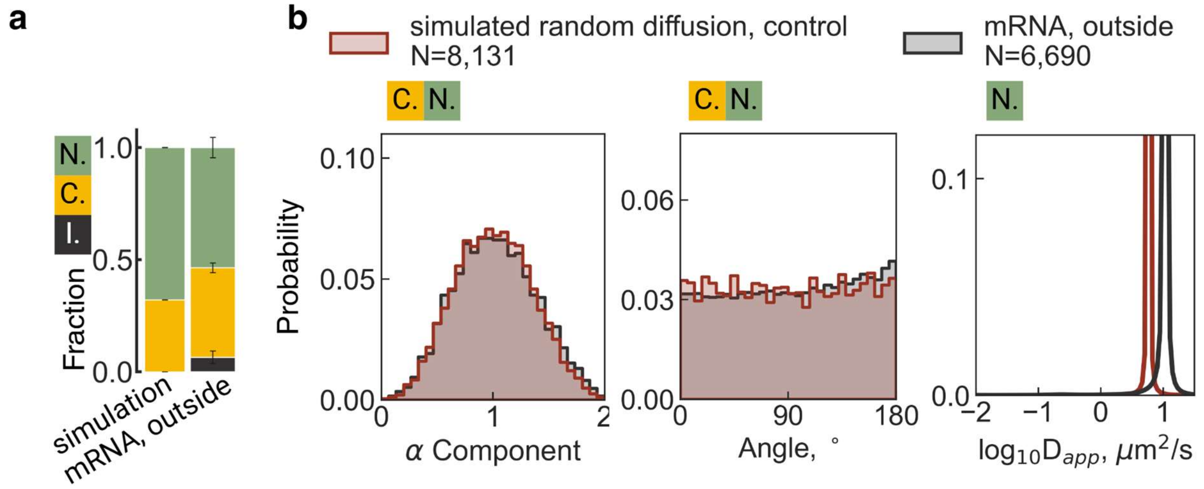


**Fig. S8 | Verification of the diffusion profiling pipeline on simulated random walk particles compared to FL mRNA in dilute FUS phase as a control for confinement threshold.**

**a |** Fractions of the three diffusion types: immobile (I.), confined (C.), and normal (N.) diffusion. Error bars for experimental data are SEM from a least three biological replicates. **b |** Distribution of anomalous component α. Distribution of angle between adjacent steps within a trajectory. Distribution of $D_{app}$ calculated using a Bayesian-based state array (SA) method[9].

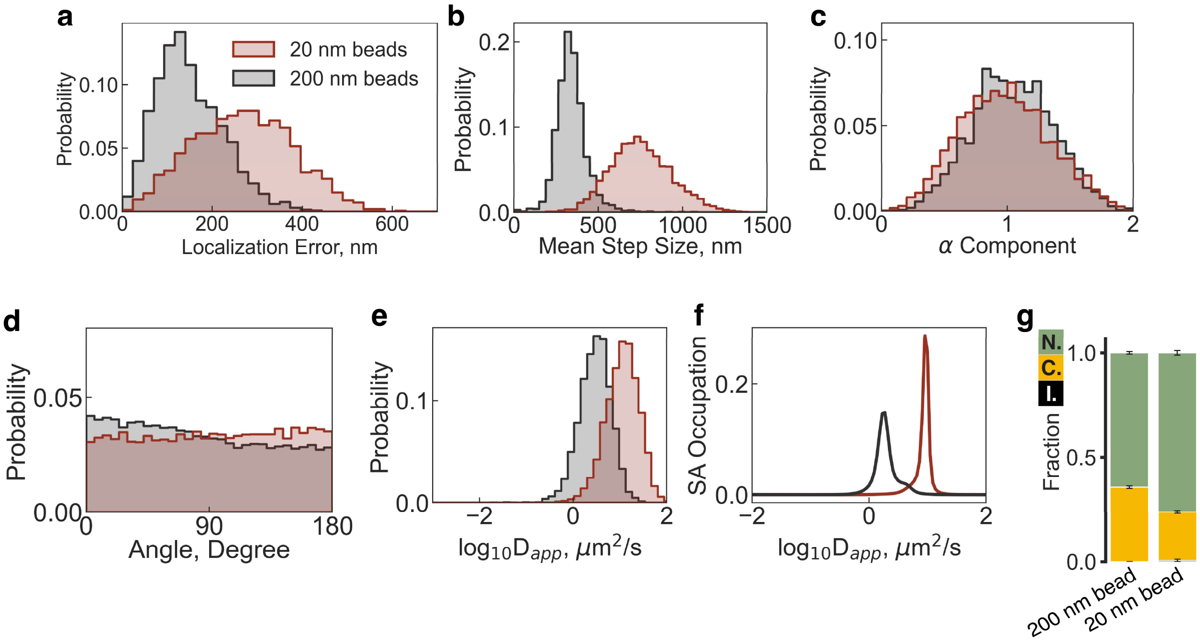


**Fig. S9 │ Verification of the diffusion profiling pipeline on 20-nm and 200-nm beads in the dilute FUS phase as a control for normal diffusion.**

**a |** Distribution of localization error extracted from an optimized form of MSD-τ fitting[7,8]. **b |** Distribution of mean step size of each trajectory. **c |** Distribution of anomalous component α. **d |** Distribution of angle between adjacent steps within a trajectory. **e |** Distribution of apparent diffusion coefficient $D_{app}$ calculated from an optimized form of MSD-τ fitting[7,8]. **f |** Distribution of $D_{app}$ calculated using a Bayesian-based state array (SA) method[9]. **g |** Fractions of the three diffusion types: immobile (I.), confined (C.), and normal (N.) diffusion. Error bars are SEM from a least three biological replicates.

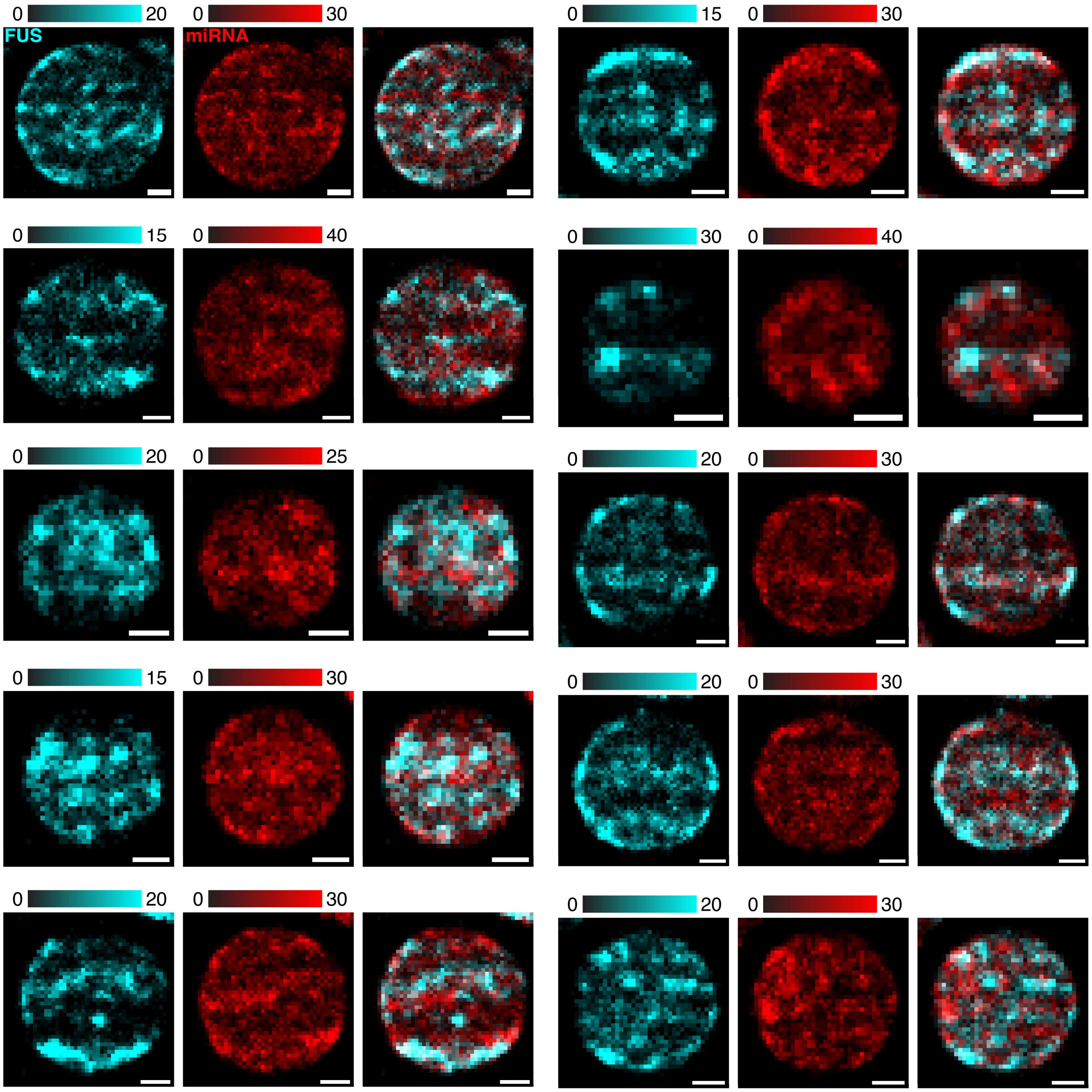


**Fig. S10 │ Representative dual-color SMT-PAINT images of FUS and miRNA-21.**

Color bars are in the unit of number of single-molecule localizations per pixel. All scale bars are 1 μm.

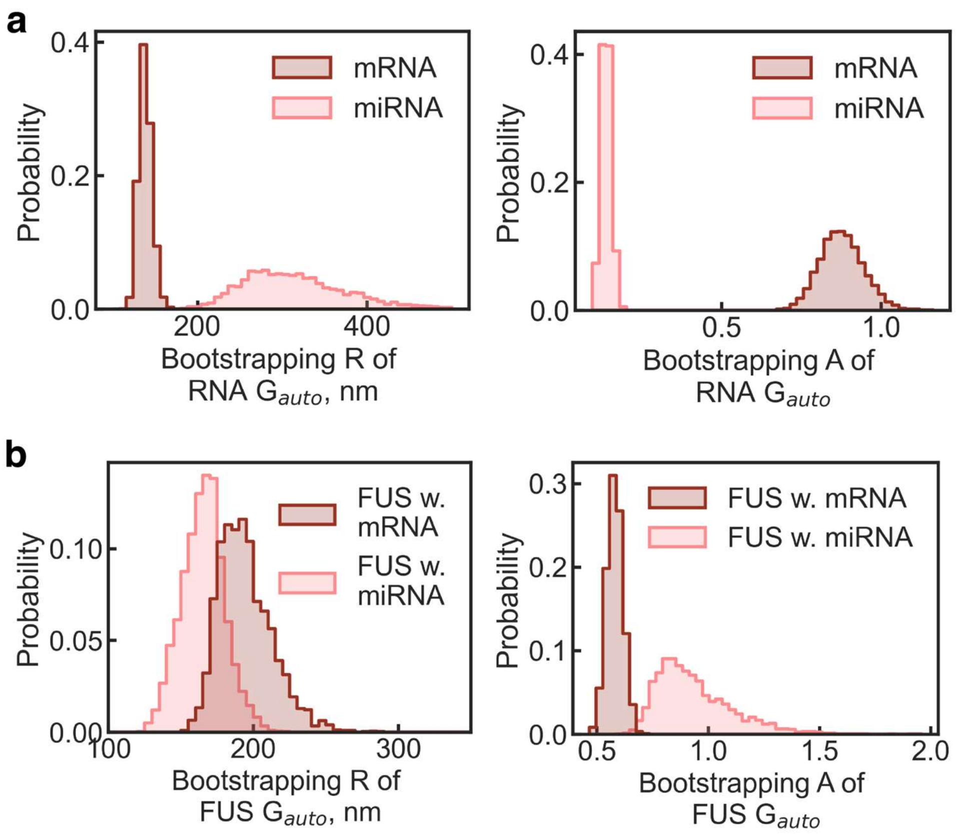


**Fig. S11 │ Nanodomain size and confinement strength estimated by bootstrapping auto-pair correlation function ($G_{auto}$)**

**a** │ Distribution of the size (R) and confinement strength (A) of RNA nanodomains extracted by parametric bootstrapping on the $G_{auto}$ of mRNA or miRNA SMT locations within each FUS condensate using an exponential decay model. **b** │ Distribution of the size (R) and confinement strength (A) of FUS nanodomain extracted by parametric bootstrapping on the $G_{auto}$ of FUS SMT locations in the presence of mRNA or miRNA using an exponential decay model.

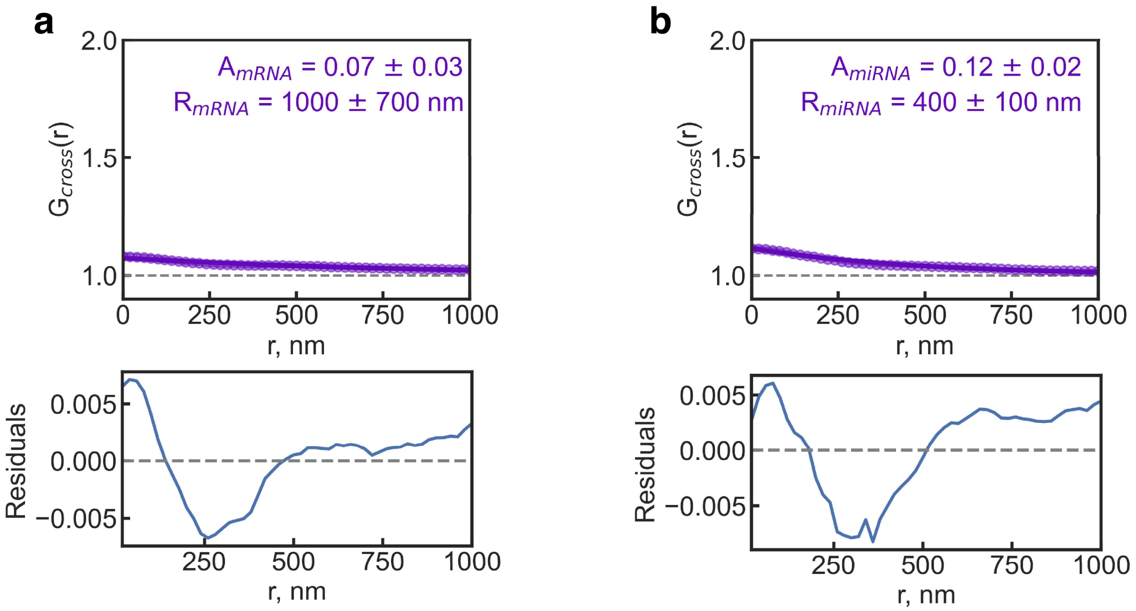


**Fig. S12 │ Marginal spatial correlation between FUS and RNA molecules revealed by a cross-pair correlation function.**

**a │** The cross-pair correlation function ($G_{cross}$) of mRNA single-molecule locations relative to FUS, with amplitude (A) and radius (R) extracted by fitting with an exponential decay with a baseline of 1. Residuals of the fit are plotted as a solid blue line. A gray dotted line in $G_{cross}$ indicates $G_{cross}=1$, suggesting a random distribution without spatial correlation between mRNA and FUS (which would be indicated by $G_{cross}>1$) or anti-correlation ($G_{cross}<1$). A gray dotted line in the residual plot indicates zero residuals. 95% confidence intervals of A and R were calculated by parametric bootstrapping. Error bars are SEM of $G_{cross}$ across different condensates, weighted by the number of SMT trajectories in each condensate. **b │** $G_{cross}$ of miRNA single-molecule locations relative to FUS with parameter estimates and fitting residuals plotted in the same way as in **a**.

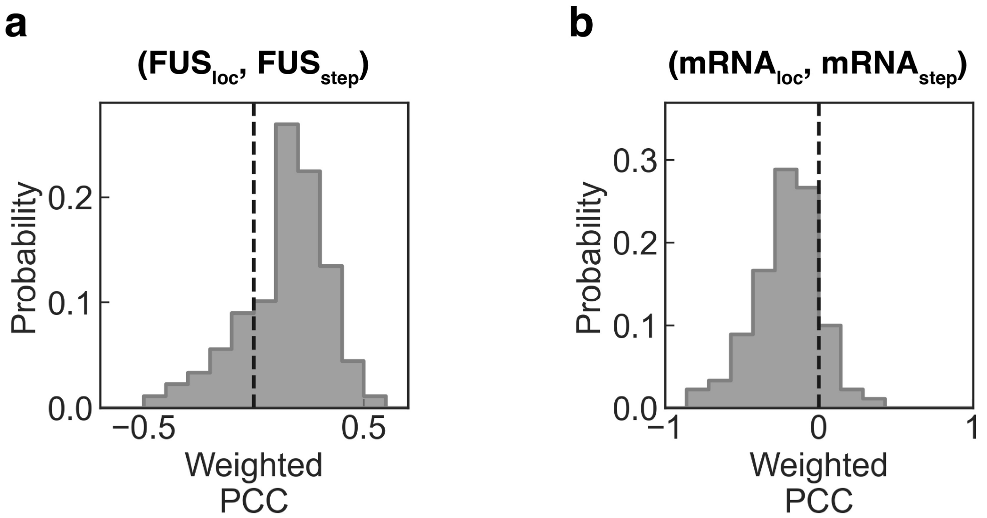


**Fig. S13 │ Penetration of fast-moving molecules into nanodomains.**

**a │** Distribution of pixel-wise Pearson Correlation Coefficient (PCC) between the SMT-PAINT images of FUS ($FUS_{loc}$) and the step size heatmap images of FUS ($FUS_{step}$), weighted by the number of SMT trajectories per pixel. A gray dotted line indicates PCC=0, separating positive correlation (PCC>0) from negative correlation (PCC<0). **b │** Distribution of pixel-wise PCC between the SMT-PAINT images of FUS ($FUS_{loc}$) and the step size heatmap images of FUS ($FUS_{step}$), weighted by the number of SMT trajectories per pixel.

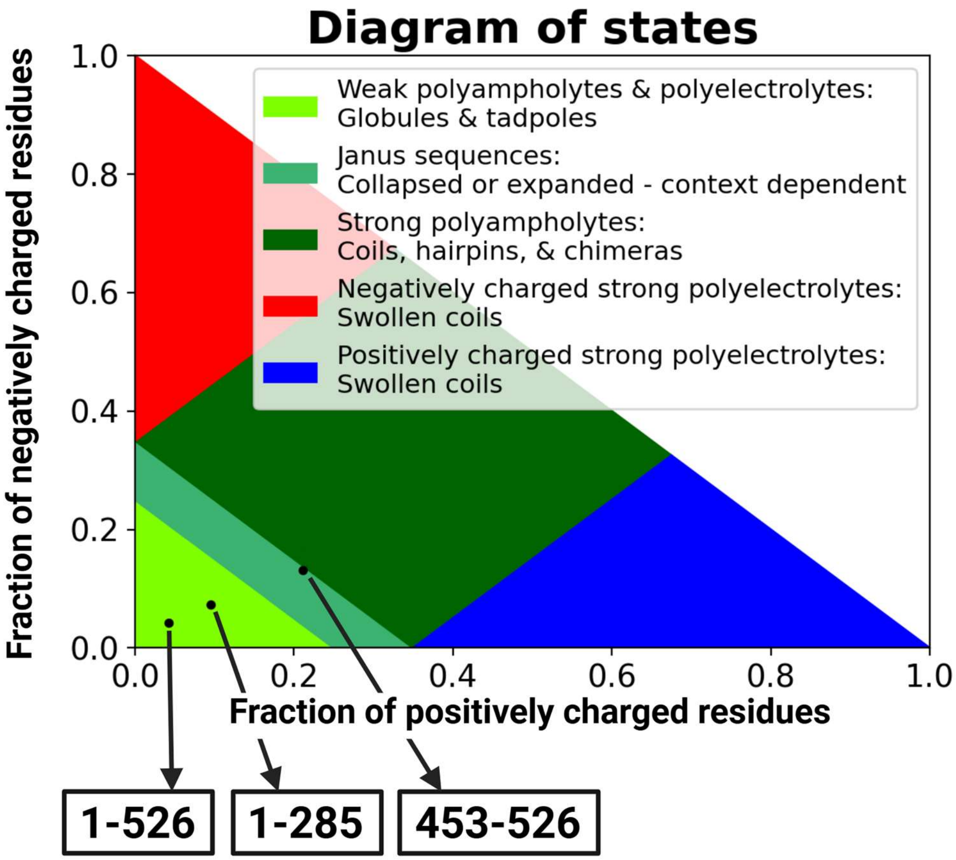


**Fig. S14 │ CIDER results for FUS**

Output from CIDER[10] for full length FUS (1-526), N-terminus IDR of FUS (1-285), and C-terminus IDR of FUS (453-526).

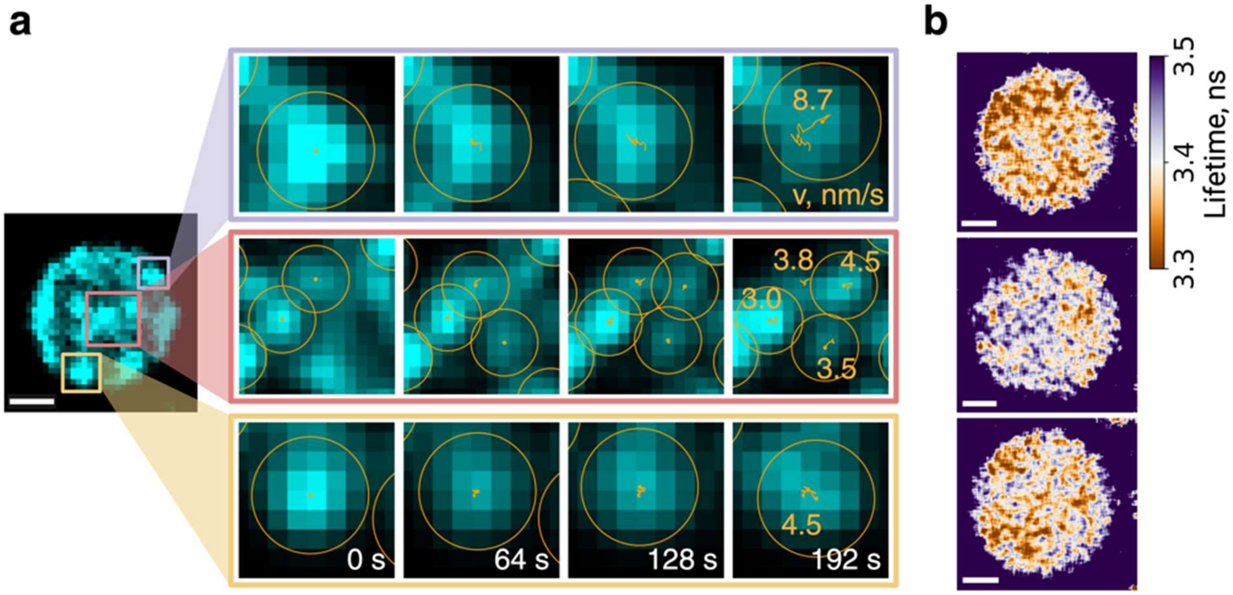


**Fig. S15 │ Nanodomains are immobile within condensates at the minutes time-scale**

**a |** Time-lapse SMT-PAINT reconstructions of nano-domains that confine FUS motions in the zoomed-in regions labeled by three different colored boxes in a SMT-PAINT image. Motions of the underlying nano-domains are revealed by orange trajectories, noted with the velocity calculated from mean step size of nanodomain trajectories. **b |** Fluorescence lifetime imaging (FLIM) of Alexa Fluor 488-labeled FUS. **c |** Cartoon and results of a random-sampling fluorescence correlation spectroscopy (FCS)-FLIM assay. The scatter plot shows the correlation between $D_{app}$ measured by FCS and lifetime by FLIM with the distributions of each on each side. The blue line shows a linear regression model fit with the shaded area being the 95% confidence interval of the fit.

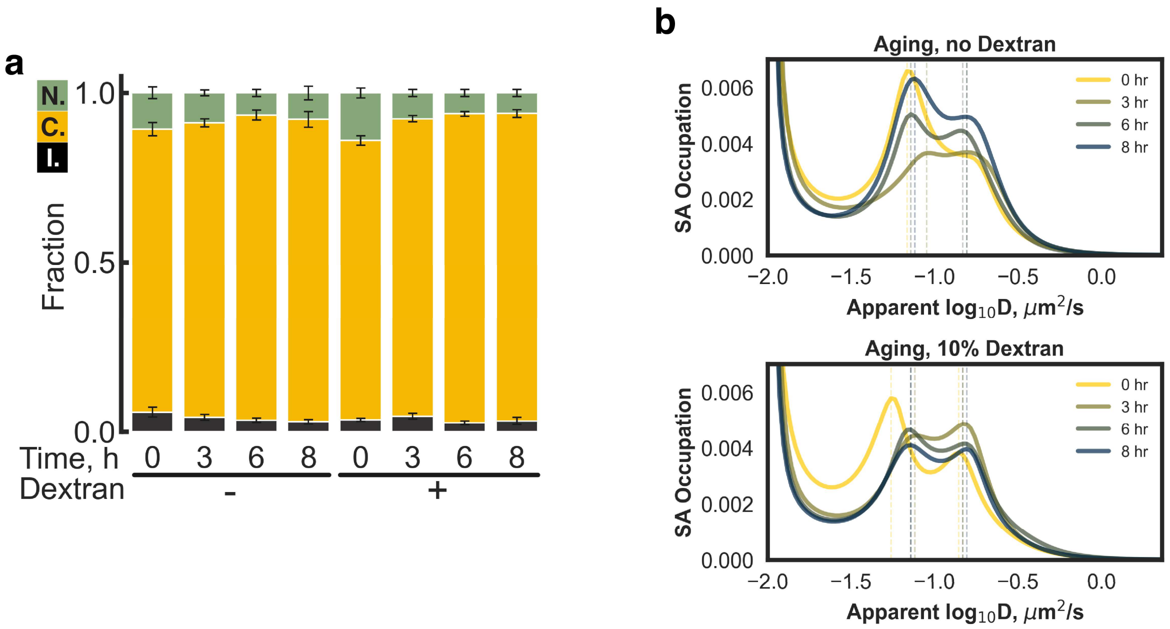


**Fig. S16 | Effect of aging and macromolecular crowding on the intra-condensate diffusion profile of mRNA molecules.**

**a** | Fractions of the three types of SMT trajectories, immobile (I.), confined (C.), and normal (N.) diffusion, of mRNA molecules within FUS condensates at varying times after reconstitution of FUS condensates with or without the crowding reagent Dextran T-500 at a final concentration of 10% (w/v). Error bars are SEM from at least three biological replicates. **b** | Distribution of $D_{app}$ calculated from normal diffusion trajectories by SA analysis, comparing the impact of aging with or without macromolecular crowding from Dextran T-500 as indicated. Vertical dotted lines indicate the peak positions.

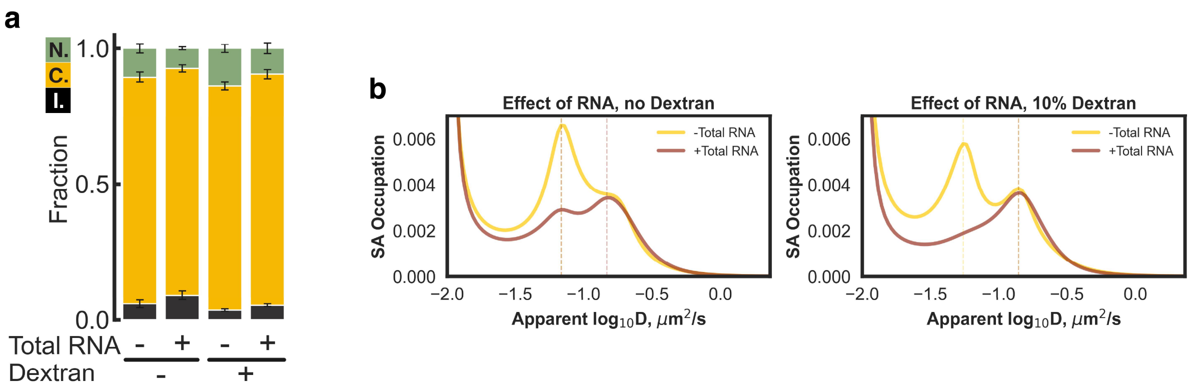


**Fig. S17 | Effect of total RNA background and macromolecular crowding on the intra-condensate diffusion profile of mRNA molecules.**

**a** | Fractions of the three types of SMT trajectories, immobile (I.), confined (C.), and normal (N.) diffusion, of mRNA molecules within FUS condensates with or without 50 ng/μL HeLa cell total RNA with or without the crowding reagent Dextran T-500 at a final concentration of 10% (w/v). Error bars are SEM from at least three biological replicates. **b** | Distribution of $D_{app}$ calculated from normal diffusion trajectories by SA analysis, comparing the impact of total RNA background with or without macromolecular crowding from Dextran T-500 as indicated. Vertical dotted lines indicate the peak positions.

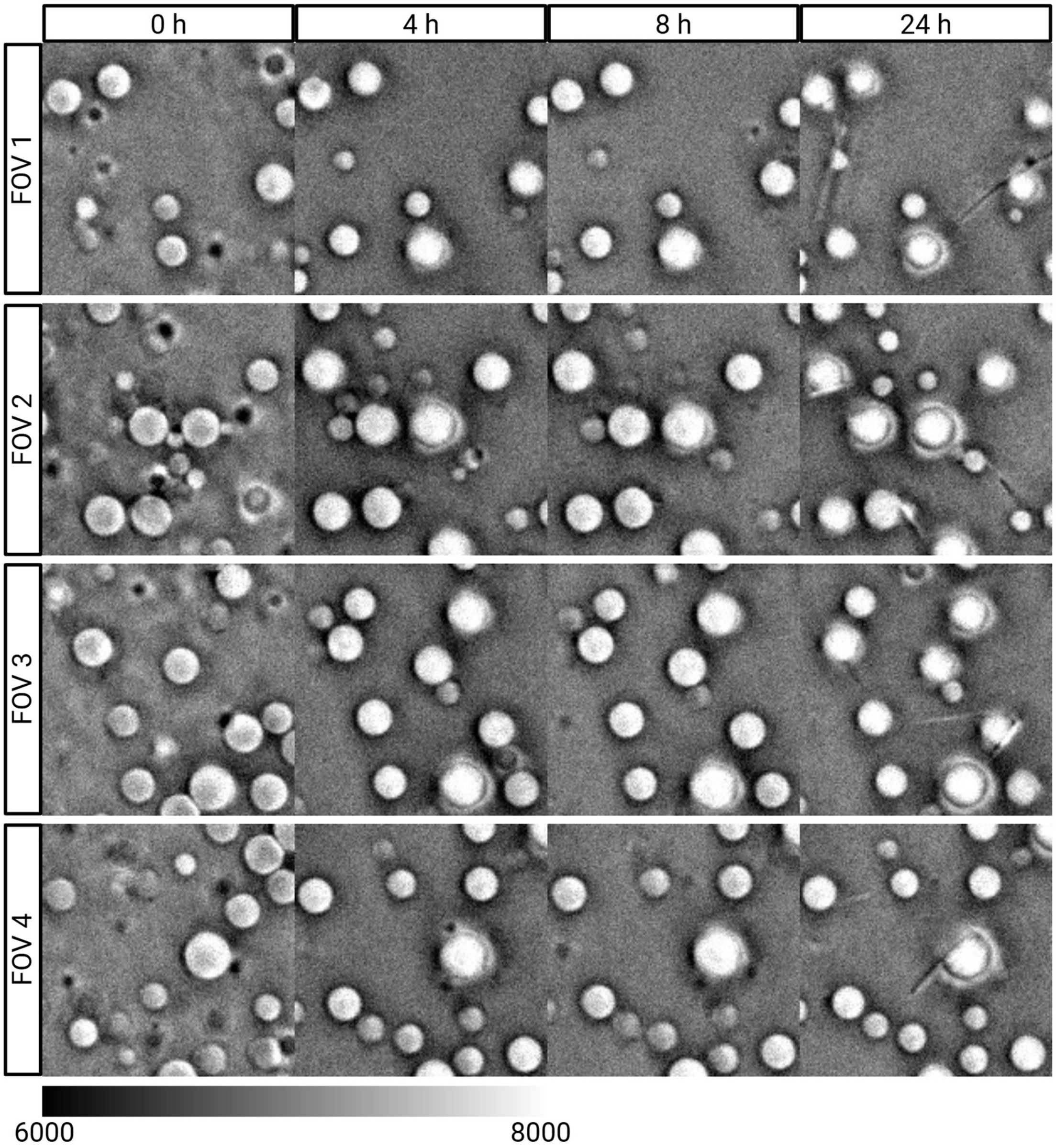


**Fig. S18 │ Bright field images of condensates in a representative FOV undergoing aging**

Four fields of view shown for FUS condensates over 24 hours. For each FOV, condensates that are liquid at time point 0h are observed to have surface fibril features at 24hrs.

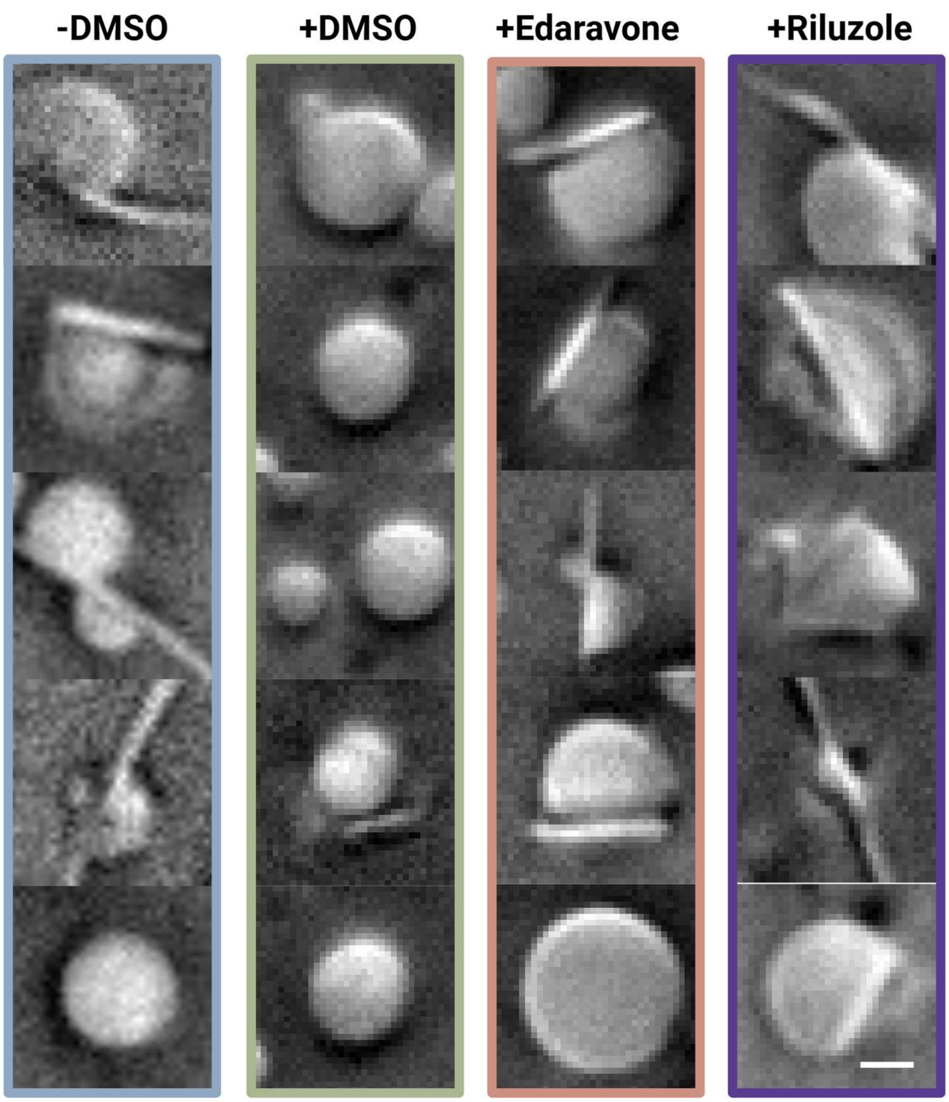


**Fig. S19 │ Small molecule drugs impact condensate aging**

FUS condensates under different small molecule drug treatment imaged in bright field at the 24-hour time point.

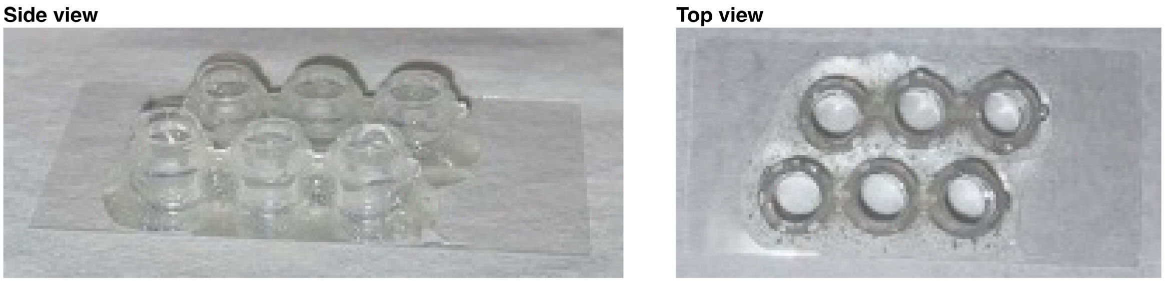


**Fig. S20 │ Representative photos of sample wells for SMT experiments.**

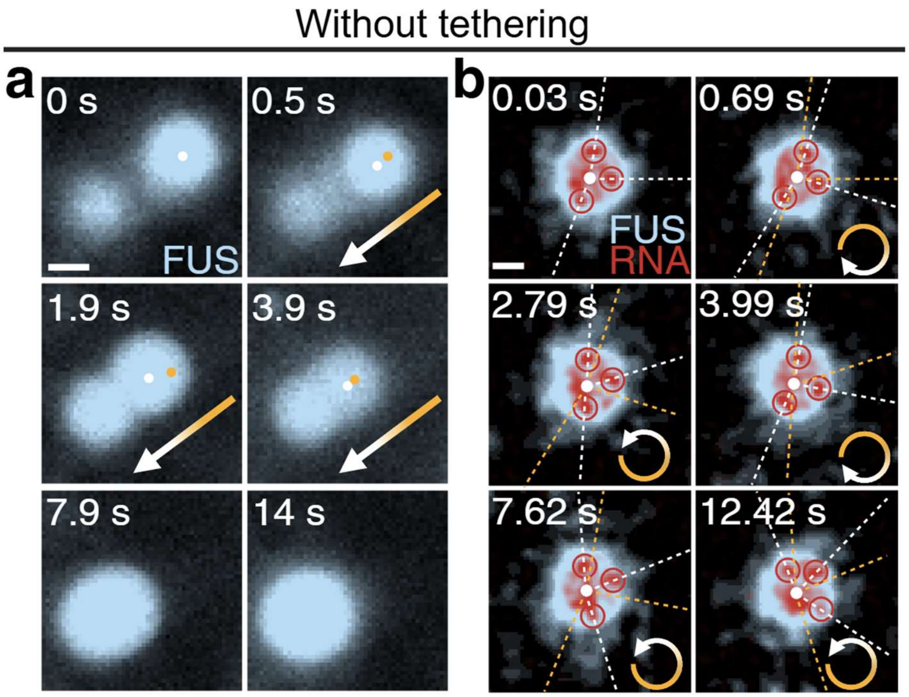


**Fig. S21 │ Translational and rotational motions of untethered condensates**

**a-b│** Condensates can undergo coalescence (**a**) and translational (arrows in **a**) or rotational (arrows in **b**) motions in the absence of FUS-biotin molecular tethers. The position of condensate (**b**) and the angular position of three RNA molecules (**b**) in the current and previous frame are shown in white and yellow respectively.

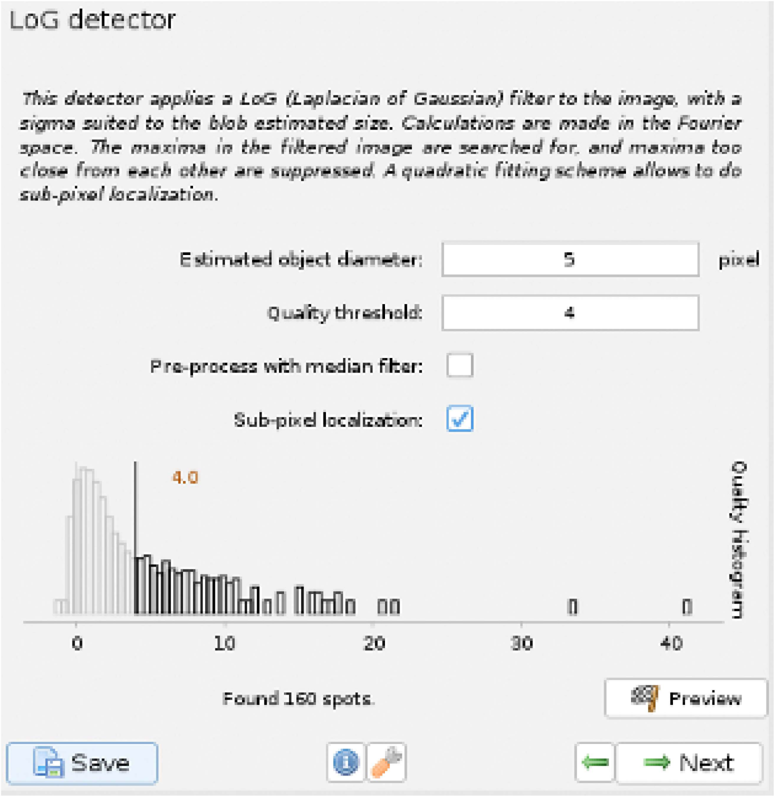


**Fig. S22 │ Representative distribution of spot quality in TrackMate, with the spot quality threshold set at the inflection point between the peak of false-positive detections and the tail of particles of interest.**

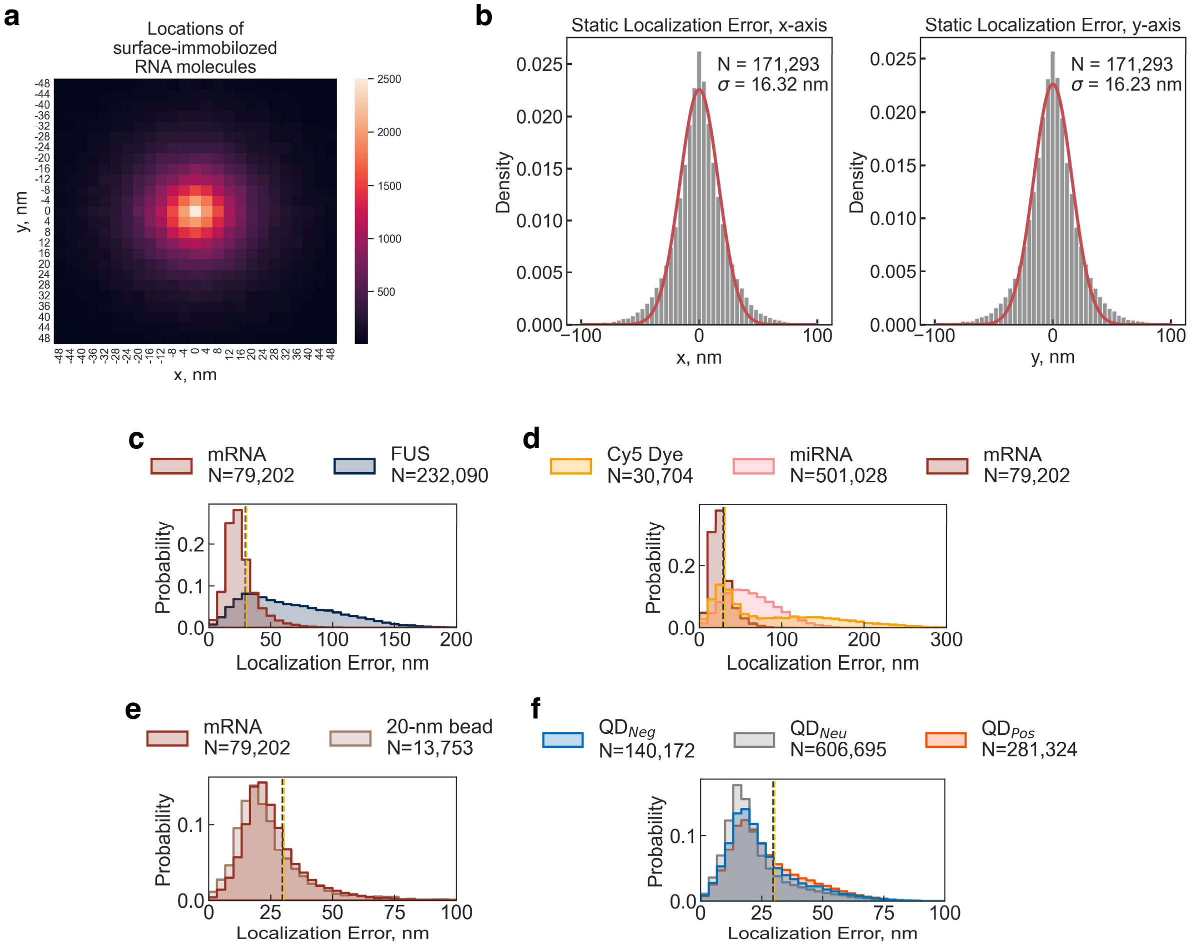


**Fig. S23 | Static and overall localization errors of intra-condensate SMT.**

**a-b** | Static localization error of Alexa Fluor 647-labeled mRNA molecules**.** To estimate static localization error, labeled mRNA molecules were immobilized onto a clean but non-passivated glass surface by non-specific surface absorption and recorded for 50 frames under the same imaging conditions used for SMT experiments. The static molecules serve as a ground truth to estimate errors introduced by fitting with a Gaussian-approximated point-spread function (PSF). For each static RNA molecules, all detected locations in a video were pooled and centered on the average of all locations from the same molecule. All centered RNA locations were pooled to estimate the deviation from the average location, assuming that the average location is the ground truth location of the static molecule. A heat map of the deviation from the centered average location is shown in **a**, while the deviation along the x or y direction is shown in **b** together with a Gaussian fit to determine the x and y localization error as the σ of the Gaussian. **c-f |** Distributions of the overall localization error calculated from an optimized form of mean square displacement (MSD)-lag time (τ) fitting[7,8] corresponding to the datasets in Fig. 2b-c (**c**), Fig. 2f-g (**d**), Fig. 2h-i (**e**), and Fig. 2j-k (**f**). Details of the fitting can be found in the *Diffusion profiling pipeline* section of the *Methods*.

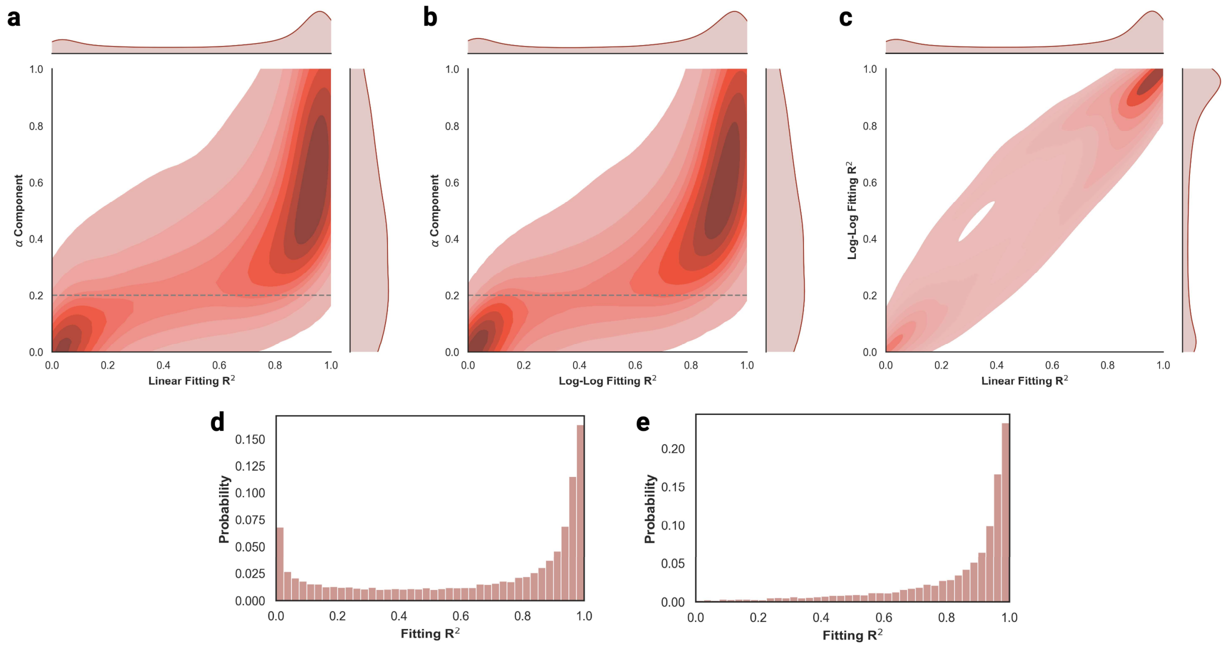


**Fig. S24 | Extreme values of the α component correlate with bad fitting to a diffusion model.**

**a |** Correlation plot of the α component versus the goodness-of-fit $R^2$ when fitting the MSD-τ curve of each SMT trajectory on a linear scale. The contour plot in the middle shows density of points with the corresponding α and $R^2$ values, while the distributions on the sides show the kernel density estimate (KDE) of distributions of α or $R^2$. The gray dotted line is a guide for the eye, denoting extremely small α values. **b |** Correlation plot of α component versus $R^2$ when fitting the MSD-τ curve of each SMT trajectory on a log-log scale. **c |** Correlation plot of $R^2$ from linear scale MSD-τ fitting versus log-log scale MSD-τ fitting. **d |** Distribution of $R^2$ for all SMT trajectories. **e |** Distribution of $R^2$ after the removal of SMT trajectories bearing extremely small α values.

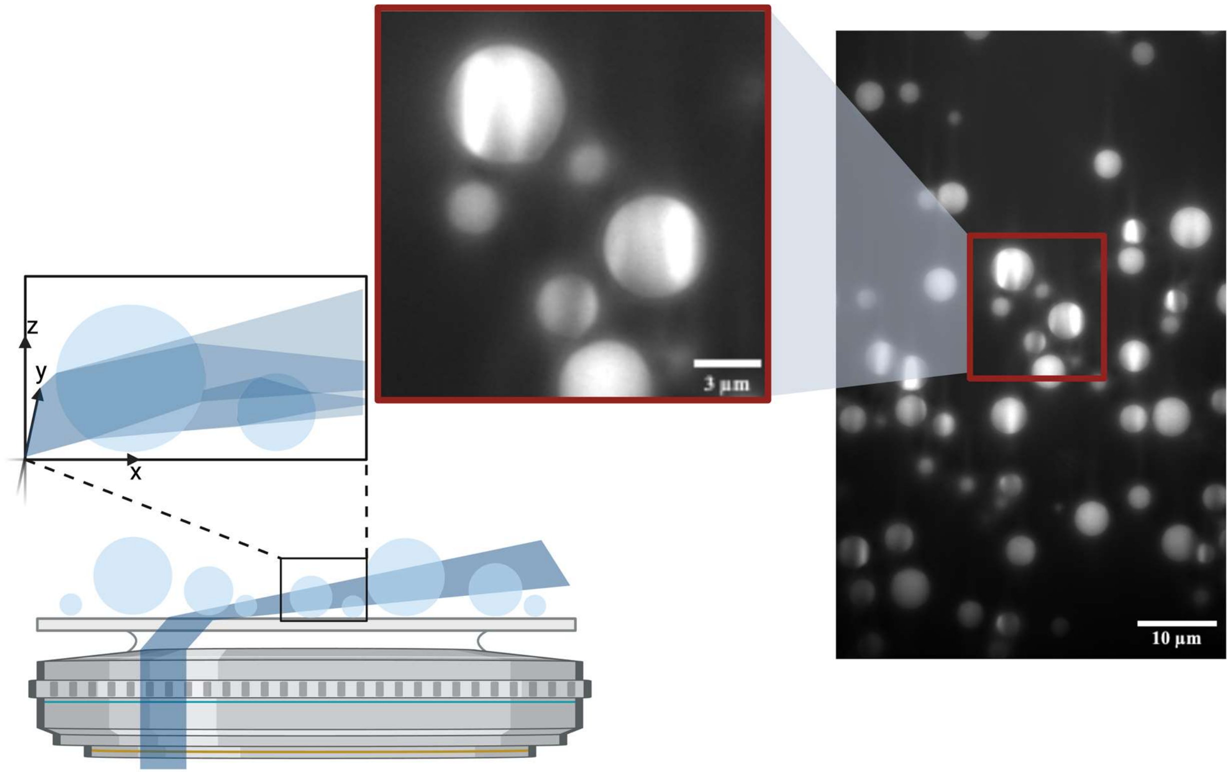


**Fig. S25 │ Lensing effect of condensates that induce stripe-like patterns upon HILO illumination of tethered FUS condensates.**

Condensates containing 10 nM Alexa Fluor 488-labeled FUS while fluorescence images were taken under 473 nm laser excitation. The spherical shape of the condensates and their distinct dielectric constant (and thus refractive index) enable them to function as lenses, refracting the incoming laser beam and focusing light onto specific regions of the condensate located behind the lens-like condensate. This lensing effect results in non-uniform illumination and thus a stripe-like pattern in the condensate behind the lens-like condensate.

### Supplemental Methods:

### Condensate Assembly and OSS usage:

Note that we keep the time between the assembly of condensates and imaging within 30 minutes, except for aging experiments where otherwise specified (Fig. 4-5), to ensure that all experimental observations reported in this study are signatures of early-stage condensates representing the dynamically assembled RNP condensates in cells. Most importantly, we added the stock solution of FUS as the last component so that phase separation was initiated only after all other components were thoroughly mixed, mimicking the assembly of condensates within intracellular milieu. A total volume of 30 $\mu$L was used for each experiment.

For intra-condensate SMT or SPT with condensate boundary detection needed (Fig. 1g, #1), 10 nM Alexa Fluor 488-labeled FUS and 50 pM of the fluorescent species for SMT or SPT, such as Cy5 free dye, Cy5-labeled miRNA-21 (Integrated DNA Technologies), Alexa Flour 647-labeled FL mRNA, 20-nm FluoSpheres Carboxylate-Modified Microspheres (Thermo Fisher), 9.5-nm CdSe/ZnS core-shell type QDs functionalized with carboxylic acid, PEG, or amine (Millipore Sigma), were mixed with phase separation buffer and an OSS composed of 2.4 mM protocatechuic acid (PCA), 24 nM protocatechuate-3,4-dioxygenase (PCD), and 2.4 mM Trolox[4] before adding stock solution of FUS. Cy5 free dye was made by neutralizing the N-hydroxysuccinimide (NHS) group on Cy5 NHS-ester (APExBIO) in 10 mM Tris-NaCl, pH 8.5 in dark at room temperature overnight. For the tracking of Cy5 free dye, single-Cy5-labeled miRNA-21, and the QDs, the PCA-PCD-Trolox OSS was omitted because these particles have low signal to noise ratio (SNR) due to fast-diffusion induced blurring, single-dye rather than multiple-dye labeling per molecule, and luminescence rather than fluorescence. For intra-condensate SMT of FUS, where condensate boundary detection was not needed because FUS single molecules in dilute phase cannot be tracked due to motion blurring by its fast diffusion (Fig. 1i, #2), 50 pM of Alexa Fluor 488-labeled FUS were mixed with phase separation buffer without OSS before adding stock solution of non-labeled full-length tag-free FUS. OSS was again omitted because Alexa Fluor 488-labeled FUS molecules have low SNR due to fast-diffusion, which we found could be further hindered by autofluorescence of high concentrations of PCA and Trolox in the OSS. However, for all trajectory reconstruction experiments, extended SMT measurements were needed for an adequate coverage of trajectories over the intra-condensate space (Fig. 1h) and thus we implemented a color-less coupled glucose oxidase and catalase (GODCAT) OSS[4] with 55 U/mL glucose oxidase, 40 mM glucose, and 290 U/mL catalase. This approach enabled us to obtain 10,000-frames long high-SNR SMT videos with much less autofluorescence than PCA and Trolox that fulfills trajectory reconstruction needs (Fig. S8). For intra-condensate SMT of FUS and RNA for dual-color trajectory reconstructions (Fig. 1i, #3), 50 pM of Alexa Fluor 488-labeled FUS and Cy5-labeled miRNA-21 or Alexa Fluor 647-labeled FL mRNA were mixed with phase separation buffer and GODCAT OSS before adding stock solution of non-labeled full-length tag-free FUS. For any experiments involved crowding or total RNA, Dextran T-500 (Millipore Sigma) was used at a final concentration of 10% (w/v) and Human HeLa Cell total RNA (TakaraBio) was used at a final concentration of 50 ng/$\mu$L.
